# Federated Multi-Site Normative Modeling using Hierarchical Bayesian Regression

**DOI:** 10.1101/2021.05.28.446120

**Authors:** Seyed Mostafa Kia, Hester Huijsdens, Saige Rutherford, Richard Dinga, Thomas Wolfers, Maarten Mennes, Ole A. Andreassen, Lars T. Westlye, Christian F. Beckmann, Andre F. Marquand

## Abstract

Clinical neuroimaging data availability has grown substantially in the last decade, providing the potential for studying heterogeneity in clinical cohorts on a previously unprecedented scale. Normative modeling is an emerging statistical tool for dissecting heterogeneity in complex brain disorders. However, its application remains technically challenging due to medical data privacy issues and difficulties in dealing with nuisance variation, such as the variability in the image acquisition process. Here, we introduce a federated probabilistic framework using hierarchical Bayesian regression (HBR) for multi-site normative modeling. The proposed method completes the life-cycle of normative modeling by providing the possibilities to learn, update, and adapt the model parameters on decentralized neuroimaging data. Our experimental results confirm the superiority of HBR in deriving more accurate normative ranges on large multi-site neuroimaging datasets compared to the current standard methods. In addition, our approach provides the possibility to recalibrate and reuse the learned model on local datasets and even on datasets with very small sample sizes. The proposed federated framework closes the technical loop for applying normative modeling across multiple sites in a decentralized manner. This will facilitate applications of normative modeling as a medical tool for screening the biological deviations in individuals affected by complex illnesses such as mental disorders.

## 1 Introduction

*Normative modeling* was recently introduced as a statistical framework for studying the biological heterogeneity of mental disorders in clinical neuroimaging cohorts (Marquand et al., 2016a). Normative modeling involves estimating the centiles of variation, *i.e.,* the normative ranges, of a biological brain measure (*e.g.*, ROI cortical thickness, ROI volume, functional connectivity) as a function of clinical covariates. This is performed via regressing the units of neuroimaging data (*e.g.*, a voxel in structural or functional MRIs) against a set of clinically relevant covariates (*e.g.*, age). Analogous to the use of ‘growth charts’ in pediatric medicine, such a mapping function provides a norm for the changes in the structure or functional dynamics of the brain across the human lifespan. Deviations of individuals from the normative range can be quantified as z-scores (Marquand et al., 2019). This approach has recently been used to dissect the heterogeneity of several mental disorders (Wolfers et al., 2018; Zabihi et al., 2019; Wolfers et al., 2020; Zabihi et al., 2020), providing compelling evidence that brain abnormalities of patients with psychiatric disorders cannot be captured in a case-control setting, *i.e.*, by average group differences between patients with a specific disorder and healthy controls. Thus, normative modeling allows us to enhance classical symptom-based diagnostics by incorporating biological and environmental factors in a principled way. Such a paradigm change will hopefully result in developing effective biological tests and individualized treatments to improve the quality of life of patients with psychiatric, neurodevelopmental, and neurodegenerative disorders (Insel and Cuthbert, 2015; Fernandes et al., 2017).

The success of normative modeling depends on the accuracy of estimating the norm and the variability around this norm for a certain brain measure (or putative biomarker) across a population. Therefore, massive data availability from a large and diverse population, extensive computational resources, and intelligent modeling techniques play pivotal roles. The advancements in data sharing standards (Miller et al., 2016) and protocols (Gorgolewski et al., 2016; Niso et al., 2018; Pernet et al., 2019) led to an exponential growth in neuroimaging data availability. Neuroimaging groups worldwide join forces in international consortia leading to clinical neuroimaging studies that are orders of magnitude larger today than a decade ago (Thompson et al., 2014; Miller et al., 2016; Casey et al., 2018). This trend has just begun, and with the recent advances in high-performance computing technologies such as grid computing, cloud computing, and GPU technologies, we now possess enough computational power to store and process these massive datasets. Furthermore, progress in artificial intelligence and machine learning over the last decades brought ubiquitous applications in healthcare. These developments are the foundations for large-scale normative modeling.

In this article, we attack the problem of estimating a *reference* normative model across a massive population using multi-center neuroimaging data and on an unprecedented scale. Developing such a reference normative model is challenged in practice by two main obstacles. First, it requires aggregating smaller neuroimaging datasets acquired at several imaging centers with different acquisition protocols and scanners. Furthermore, the data is often processed using various preprocessing pipelines and toolboxes, each of which leaves its signature on the final derived statistics (Fortin et al., 2018), referred to as *site-effects*. Site-effects introduce artefactual variability in data that confound the derived deviations in normative modeling (Marquand et al., 2019). In this way, the practical application of normative modeling as a medical tool is limited as the data collected at different centers may have different characteristics. Thus, it is essential to develop effective methods that can effectively deal with these confounds. The second barrier toward developing a reference normative model and deploying it as a medical tool at local clinical centers is data privacy (Poline et al., 2012; Rieke et al., 2020). Clinical data are always subject to privacy regulations and cannot be distributed freely without acquiring appropriate consent. This fact challenges the centralized model estimation in which the estimation algorithm requires access to whole data at once. Therefore, it is essential to decentralize the model estimation phase by developing a federated learning (McMahan et al., 2017; Yang et al., 2019b; Rieke et al., 2020; Li et al., 2020) approach for normative modeling.

In this work, we address these problems by extending our previous work on multi-site normative modeling (Kia et al., 2020). We propose hierarchical Bayesian regression (HBR) (Gelman et al., 2013) for probabilistic modeling of group-effects (including batch-effects) in neuroimaging data. The proposed model is hierarchical in the sense that it imposes shared prior distributions over site-specific model parameters. Our method has several appealing features: i) it is fully probabilistic, thus, it is well-suited to normative modeling as it provides estimations of both phenomenological variability in data and epistemological uncertainty in the model (Cox, 2006); ii) it preserves all sources of variation in the data while ensuring unwanted site-related variability is not reflected in the output statistics; iii) it is highly flexible and accommodates different modeling choices (*e.g.*, non-linear effects or heteroscedastic noise); iv) it provides the possibility of transferring information from a reference model when re-calibrating the normative model to even small data from new sites, and v) it provides the possibility of model estimation/calibration on decentralized data.

This work extends our previous conference publication in Kia et al. (2020) from both methodological and experimental aspects. From the methodological point of view, here we exploit the generative nature of the HBR model to estimate/update model parameters on decentralized data in the context of the life-cycle of normative models. Furthermore, we scaled up the size of our experimental data to 16 datasets of 37126 participants from 79 scanners. Our experimental results replicate and extend the results in Kia et al. (2020) demonstrating the superiority of HBR in estimating the predictive posterior distribution compared to the prevalent ComBat data harmonization approach (Fortin et al., 2017) and trivial pooling. We also show the benefit of transferring prior knowledge in scenarios where data are extremely sparse, even down to a few data points per scan site which is still encountered in clinical applications.

## 2 Materials and Methods

In this section, we first sketch the life-cycle of the normative model as a clinical tool. We discuss the components involved in this life-cycle of normative modeling and their technical requirements. Then, we formally show how the hierarchical Bayesian framework is used to fulfill those requirements.

### 2.1 Normative Modeling: The Life-Cycle

The life-cycle of a normative model, Fig. 1, is comprised of two main components i) model development, and ii) model deployment. The model development refers to the process of estimating the parameters of the reference model on a large set of multi-center data. The model deployment refers to the process of adapting the parameters of the reference model to local data, *e.g.*, at local hospitals or research centers.

**Figure 1:**
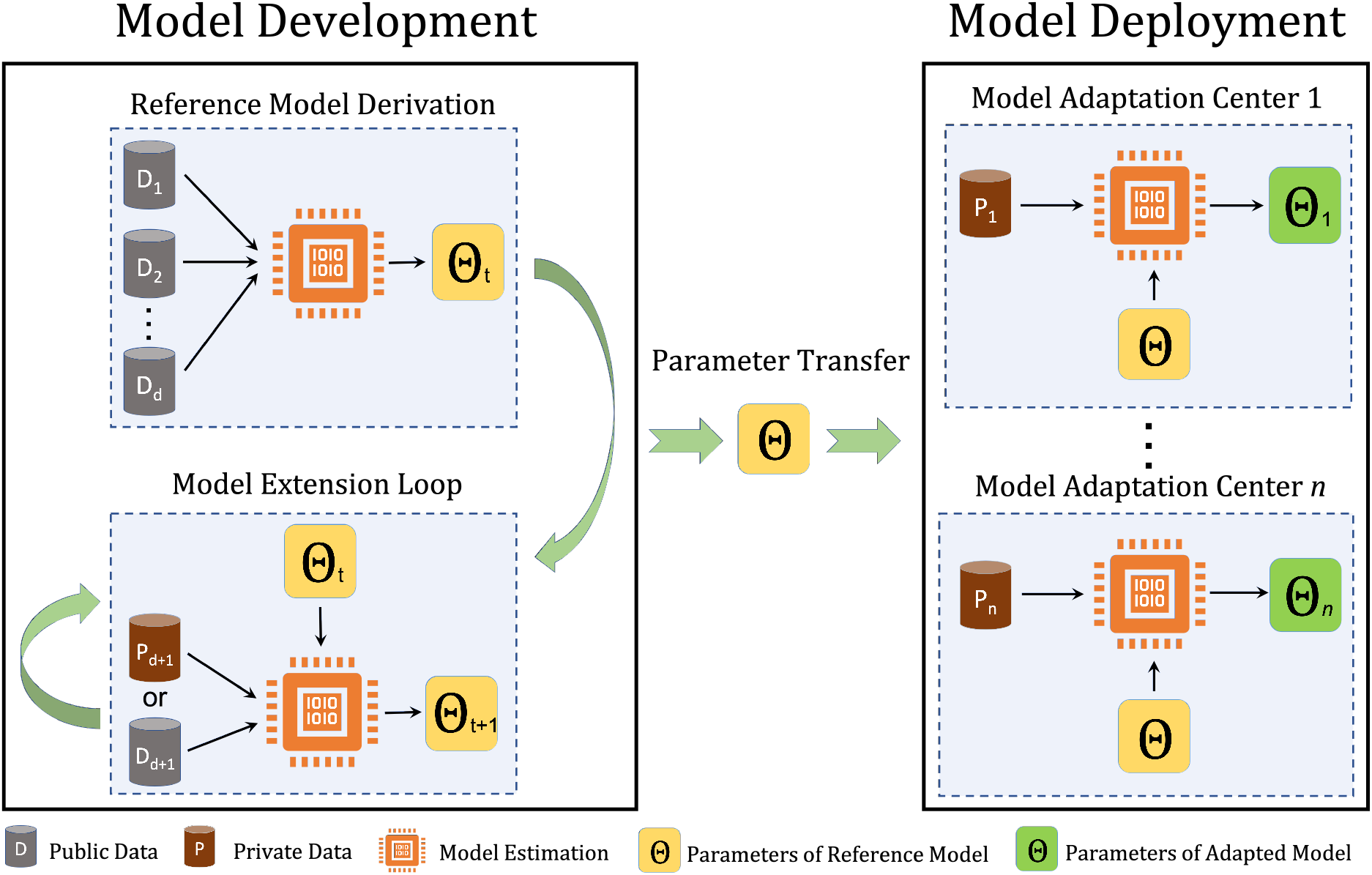
Model development and model deployment in the normative model life-cycle. In the model development phase, the parameters of the reference model are estimated on *d* datasets (*D*_1_*, D*_2_*, …, D_d_*). The model extension loop provides the possibility of model development on decentralized data at time point *t*. In the model deployment phase, the parameters of the reference model are adapted to local data at hospitals or research centers.

Considering the real-world limitations in developing and deploying a reference normative model (multi-site data and data privacy/access issues), the modeling approach that is employed for estimating the parameters of the normative model must have four vital features:

1. It should be able to deal with site-effects;
2. In the development phase, it should have the possibility of updating the parameters of the reference model over time and when new datasets are available, without requiring access to the full primary dataset. We refer to this process as model extension;
3. It should apply to both centralized and decentralized data. The centralized data refers to the scenario in which all training data are available for model estimation. In the decentralized case, the data are distributed across different centers, and data sharing and transfer are not possible due to privacy issues;
4. In the deployment phase, it should provide a mechanism for adapting the parameters of the reference model to novel data at the deployment centers (*e.g.*, local hospitals). It is crucial to emphasize that the initial data used for estimating the reference model might not be available during the adaptation process. Therefore, the knowledge transform must be performed using a parameter transfer learning approach (Pan and Yang, 2009). We refer to this process as model adaptation.

We emphasize that the model adaptation is different from the model extension process. The model extension is used during the reference model development in which we aim to derive a larger model from a smaller one. Whilst model adaptation is used in the model deployment where we aim to distill a smaller model from a reference model on local data.

In the remaining text of this section, we present practical solutions for implementing these features. To this end, we first formally define the normative modeling procedure.

### 2.2 Normative Modeling: The formal Definition

Let **X** ∈ ℝ^*n*×*p*^ represent a matrix of *p* clinical covariates for *n* participants. We denote the corresponding neuroimaging measures at each measurement unit (*e.g.*, a voxel) by **y** ∈ ℝ^*n*^. Assuming a Gaussian distribution over each neuroimaging measure, i.e., 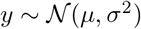, in normative modeling we are interested in finding a parametric or non-parametric form for *μ* and *σ* given the covariates in **X**. Then, for example, *μ* ± 1.96*σ* forms the 95% percentile for the normative range of **y**. To estimate *μ* and *σ*, we parametrize them respectively on *f_μ_*(**X**, *θ_μ_*) and 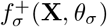, where *θ_μ_* and *θ_σ_* are the parameters of *f_μ_* and 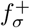. Here, 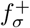 is a non-negative function that estimates the standard deviation of heteroscedastic noise in data. The homoscedastic formulation is a specific case where *σ* is independent of **X**. The non-negativity of 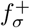 can be enforced for example using the *softplus* function 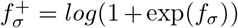 (Tanno et al., 2017; Lakshminarayanan et al., 2017; Patro et al., 2019).

In the normative modeling context, the deviations of samples from the normative range are quantified as z-scores (Marquand et al., 2016a):

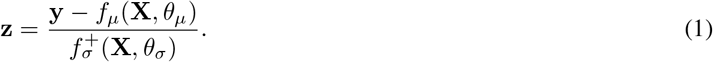

As discussed, to close the application loop for normative modeling, the model must accommodate multi-site data. We discuss classical strategies for multi-site neuroimaging data modeling in the next section.

### 2.3 Multi-Site Normative Modeling

Let 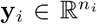 denote neuroimaging measures for *n_i_* participants in the *i*th group, *i* ∈ {1, …, *m*}, of data and we have 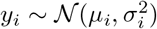. Here, a group refers to any non-ordinal categorical variable such as a batch-effect (that causes unwanted and non-biological variation in data) or other biologically relevant variables such as sex or ethnicity. In this article, since the focus is on multi-site normative modeling, we use the term ‘batch’ to refer to each group (otherwise mentioned) where each batch refers to data that are collected at different imaging sites. However, our formulations are general for application on other possible batch-effects (*e.g.*, processing software version) or biologically relevant group-effects (*e.g.*, sex and ethnicity).

Traditionally, there are four possible strategies for normative modeling on multi-site data. In the following, we explain the theoretical and practical limitations of these approaches in the life-cycle of normative modeling.

#### 2.3.1 Naive Pooling

Naive pooling is a variation of the complete pooling scenario (see Fig. 3) in which the batch-effects in data are ignored by assuming that data in different batches are coming from the same distribution, *i.e.*, 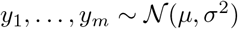 and we have:

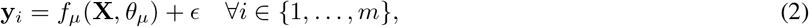

where *ϵ* is zero-mean error with standard deviation 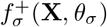. Even though the naive pooling approach provides a simple solution to benefit from a larger sample size, the oversimplifying assumption on identical data distributions restricts its usage in normative modeling because batch-effects are reflected on the resulting statistics in Eq. 1.

**Figure 2:**
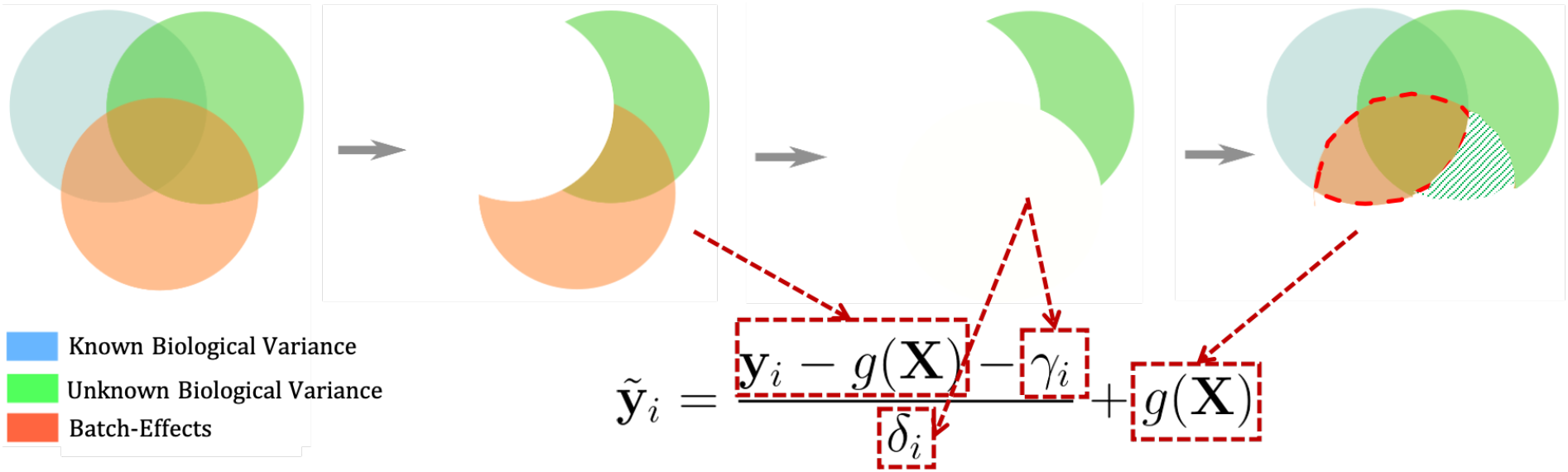
A schematic illustration of steps in data harmonization using ComBat. ComBat, firstly, removes the biologically relevant variance, which is defined in the design matrix **X**, from data. Then, it removes the additive (*γ_i_*) and multiplicative (*δ_i_*) batch-effects (here, to simplify the visualization we assume *δ_i_* = 1, ∀*i* ∈ [1, …, *m*]); and in the end, it returns the biologically-relevant variance to the residuals. This process results in removing the shared variance between the batch-effect and unknown biological sources (the green striped area) while part of the batch-effect that correlates with known sources of variance is preserved in the harmonized data (the intersection between blue and red circles).

**Figure 3:**
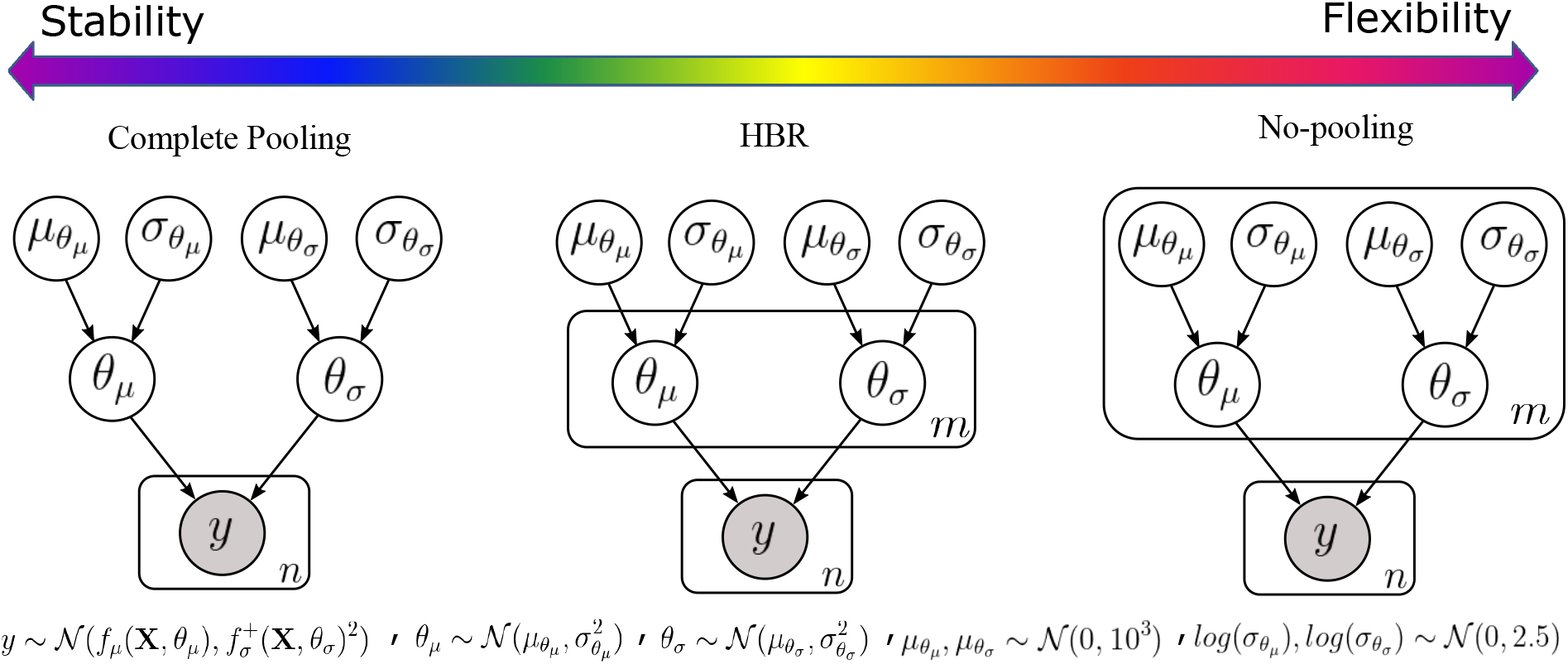
Graphical models of complete pooling, partial pooling via HBR, and no-pooling. The solutions for handling the site-effects form a spectrum in the model stability-flexibility space. At the stability end of the spectrum, we have the complete pooling solution. In the complete pooling scenario, the model learns the same set of parameters and hyperparameters on big data. At the flexibility end of the spectrum stands the no-pooling approach, where a large set of parameters and hyperparameters are estimated for each site, however, it does not benefit from the richness of big data. Therefore, its parameters can be unstable for sites with small sample size. The HBR lies in the middle of the spectrum, thus, it brings the best of two worlds together. In HBR, similar to no-pooling, we allow the model to learn different sets of parameters for data from multiple sites. At the same time, similar to complete pooling, the model has a fixed set of hyperparameters. Here, hyperparameters play the role of a joint prior over the parameters. They perform as a regularizer and prevent the model from overfitting on small batches.

#### 2.3.2 Pooling after Data Harmonization

In this approach, data are harmonized for batch-effects before pooling. Data harmonization overcomes the limitation of the naive pooling approach by adjusting the location and scale of the data for batch-effects. Hence, unlike naive pooling, assuming identical data distribution across batches is no longer a restrictive issue. Adopted from genomics, ComBat (Johnson et al., 2006) is a popular method for harmonizing neuroimaging data. ComBat uses an empirical Bayes method for adjusting additive and multiplicative batch-effects in data. It has shown great potential in harmonizing different neuroimaging data modalities, including diffusion tensor imaging (Fortin et al., 2017), cortical thickness (Fortin et al., 2018; Beer et al., 2020), and structural/functional MRI (Nielson et al., 2018; Yamashita et al., 2019; Pomponio et al., 2020).

ComBat removes additive and multiplicative batch-effects while preserving the signal of interest in data:

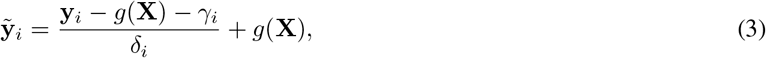

where 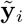 is harmonized data that is expected to be homogeneous across batches; *γ_i_* and *δ_i_* are respectively the additive and multiplicative batch-effects. Here, *g*(**X**) is a linear or non-linear (Pomponio et al., 2020) function that preserves the signal of interest as specified in the design matrix **X**. After harmonization, Eq. 2 can be used for modeling the pooled data.

However, ComBat (and in general data harmonization) has two problematic theoretical shortcomings. We refer to these problems as theoretical because depending on the covariance structure of data they might or might not occur in practice. Fig. 2 schematically illustrates these two theoretical problems when harmonizing neuroimaging data using ComBat. The signal variance in neuroimaging recordings consists of three main components: 1) the known variance in data that we *a priori* are aware of their sources, thus, we can account for them when harmonizing data (using *g*(**X**)). Examples of these sources are age and sex of participants; 2) the unknown variance in data that might have some clinical relevance, however, we have no prior information about them. An example is subtypes in psychiatric disorders. And 3) the last component is the nuisance variance related to batch-effects in data, such as site-effects that we intend to remove from data. We emphasize that these three sources of variation often correlate with each other. For example, it is very common that the participants from a certain dataset have a specific age range. In this case, the site-effect correlates with age. These shared variances are depicted as overlaps between the circles in Fig. 2.

ComBat removes all variance associated with batch-effects and preserves *a priori* known sources of variation in data (which are accounted for in the design matrix **X**) and unknown sources of variation that are not correlated with batch-effects. In other words, it is necessary to specify in advance which shared variation should be retained. This requirement is restrictive especially when we are interested in an exploratory analysis of unknown biologically relevant factors (see our simulation experiments in the supplementary). An illustrative example is stratifying psychiatric disorders into subtypes (Marquand et al., 2016b). Since subtypes are unknown in advance, their biological correlates in brain images can be removed or corrupted in the data harmonization process (the green striped area in Fig. 2). Moreover, in many cases, clinical covariates (such as age) strongly correlate with batch-effects, thus, preserving the age effect may result in a partial presence of unwanted batch-effects in the harmonized data (the intersection between red and blue circles in Fig. 2).

Data harmonization via ComBat also suffers from a practical issue when adopted in the normative modeling life-cycle. ComBat requires access to data from all sites at the training time to compute the parameters *g*(**X**), and the variance of the noise. This drawback is problematic for updating the model parameters, model estimation on decentralized data, and model adaptation to local data. Because in these scenarios, we may not have access to all data due to data anonymity concerns or a lack of ethical permission for data sharing (Poline et al., 2012). Recently, Pomponio et al. (2020) presented an ad-hoc solution to this problem in a web application. This method is based on demeaning and rescaling the data from a new site using respectively the mean and standard deviation of residuals. However, the effectiveness of this approach in removing the batch-effects while preserving the signal of interest remained unexplored and needs further empirical evaluations.

#### 2.3.3 Pooling with Batch-Effects as Fixed-Effect

In this setting, the batch-effects are directly used as covariates (in the design matrix *X*) in Eq. 2. While effective in removing the batch-effects, this method suffers from the same theoretical and practical limitations of data harmonization. It regresses out the batch-effects, thus, part of the unknown but informative variance of interest in data that are correlated with batch-effects. Model adaptation and extension procedures are also restricted in this setting because it requires full data availability. In other words, since all sites need to be encoded in the design matrix at training time, it is difficult to deploy pre-trained models to new sites.

#### 2.3.4 No-Pooling

In the no-pooling scenario, we assume that the data in each batch are drawn from different distributions. Hence, a separate and independent set of model parameters are estimated for each batch (see Fig. 3):

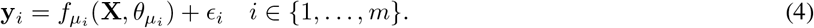

No-pooling is immune to the theoretical problems of fixed-effect pooling and harmonization because the batch-effects are not directly removed from the data. However, it cannot take full advantage of the richness of big data. It is prone to overfitting, especially when 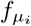 and 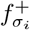 are complex functions and the number of samples in each batch is small; This may result in spurious and inconsistent estimations of parameters of the model across different batches (see results in Sec. 3.1 and 3.4.1).

### 2.4 A solution: Partial-Pooling using Hierarchical Bayesian Regression

To overcome the aforementioned shortcomings, we propose a partial pooling approach based on hierarchical Bayesian regression (HBR) as a possible solution for completing the life-cycle of normative modeling.

HBR is a natural choice in modeling different levels of variation in data (Gelman et al., 2013). In HBR, the structural dependencies between variables are incorporated in the modeling process by coupling them via a shared prior distribution. To adopt HBR for multi-site normative modeling, we assume 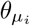 and 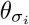 in Eq. 4 (that govern the data generating process for each batch **y**_*i*_) are coming exchangeably from the same prior distribution, *i.e.*, 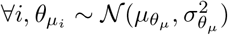 and 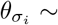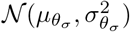 (see Fig. 3). Such a joint prior acts like a regularizer over parameters of the model and prevents it from overfitting on small batches. Given our limited prior knowledge about the distribution of parameters in the development phase and to ensure unbiased parameter estimation even for outlier sites, we use a wide Gaussian distribution as a weakly-informative hyperprior over parameters of the priors 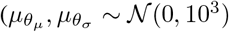 and 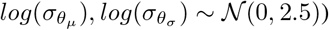.

HBR allows for a reasonable compromise between the complete pooling and no-pooling scenarios in the stability-flexibility spectrum as it combines all models in Eq. 4 into a single model that benefits from the wealth of big data, thus results in more stable models. At the same time, like no-pooling, it estimates a separate set of parameters, thus different *f_μ_* for and 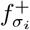 for each batch (or group). Then in the normative modeling setting, the deviations (z-statistics) for the *i*th batch are computed as follows:

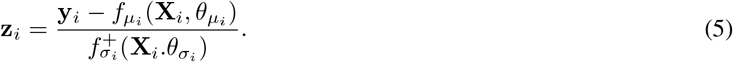

By using separate *f_μ_* and 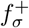 across batches, the z-statistics are respectively compensated for the additive and multiplicative batch-effects. Therefore, they are immune to batch-effects (see results in Sec. 3.1). In addition, unlike harmonization, HBR does not directly remove batch-related variability from data, thus, it preserves unknown sources of biological variations that correlate with batch-effects in data (see our simulation experiments in the supplementary).

HBR also presents several appealing features that make it the first choice for sustainable normative modeling. The generative nature of the model and shared prior distribution over parameters facilitate the model extension and adaptation, especially when dealing with decentralized data. Hence, it fulfills the technical requirements of normative modeling life-cycle (see Sec. 2.1). Furthermore, HBR provides the possibility to account for more than one group-effect and as a result more than one batch-effects in data. This is a favorable feature when we intend to simultaneously deal with several batch-effects in data (for example variability in both scanners and preprocessing software). In addition, it provides the possibility to include other informative group-effects (such as sex and ethnicity) in the hierarchical modeling process of the HBR.

#### 2.4.1 Model Extension using HBR

Considering the Bayesian nature of the HBR, once the parameters and hyperparameters of the model for a specific brain measure **y**_*i*_ are inferred, we can use the generative nature of the model to simulate synthetic neuroimaging measures **ŷ**_*i*_ by sampling from the posterior predictive distribution of the model. In the normative modeling context, each generated sample represents the data for a single healthy participant. We exploit this property to implement the extension loop in the model development process. The model extension loop in Fig. 1 can be expanded to a repetitive process of data generation and model estimation as illustrated in Fig. 4 and in the experiments below (see the model extension experimental setting in Sec. 2.6.2). Here, we assume that we have access to the data from a single dataset at stage *i* of the model estimation. To estimate the model parameters at stage *i*, the synthetic data generated for 1, …, *i* − 1 stages are used to set up the complete dataset for parameter estimation.

**Figure 4:**
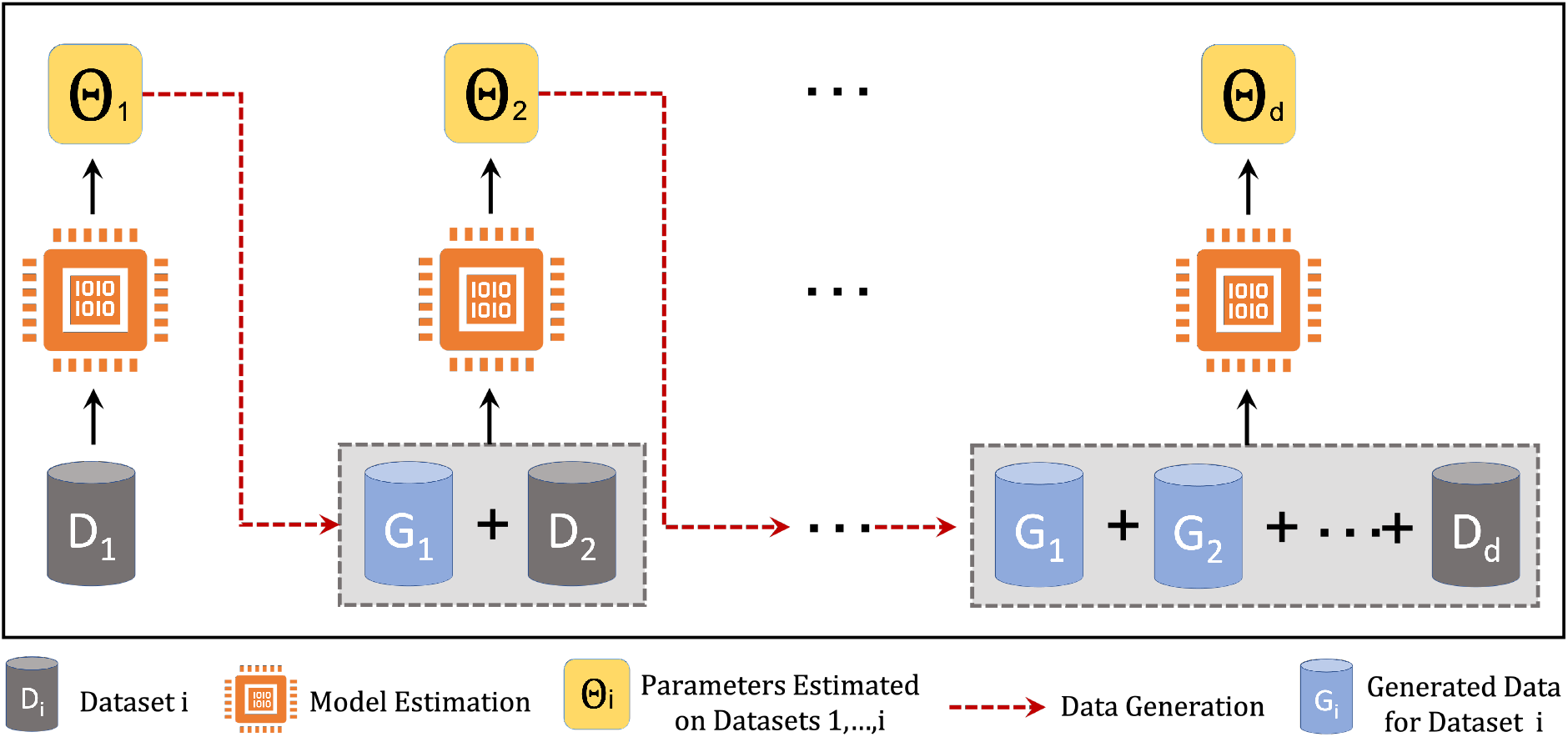
Model extension loop in multi-site normative modeling using HBR. The synthetic data generated for stages 1, …, *i* − 1 are used to estimate the model parameters at stage *i*. Model extension prodes the possibility of updating the model parameters over time, and parameter estimation on decentralized data.

In this scheme, if each stage is defined as a time interval, then the model extension loop can be used to update the model parameters over time and when new datasets are available. On the other hand, if each stage is defined as the geographical data distribution across data centers, then the model expansion loop can be used to train a reference normative model on decentralized data. These characteristics are crucial to maintaining the life-cycle of normative modeling.

#### 2.4.2 Model Adaptation by Transferring Parameters

Importantly, HBR also provides the possibility to transfer the knowledge inferred about the distribution of hyperparameters from a primary set of observed data **y** (in the model development process) to secondary datasets from new sites **y**^∗^ when deploying model at local centers. To achieve this, we propose to use posterior distributions of hyperparameters of the reference normative model, *i.e.*, 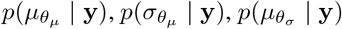, and 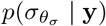, as *informative* hyperpriors for the secondary model. Informative hyperpriors enable us to incorporate pre-existing evidence when re-inferring the model on new data rather than ignoring it by using non-informative or weakly informative hyperpriors. This is also a critical feature for model portability because it enables effective model adaptation without the need to have access to the primary set of data.

### 2.5 Anomaly Detection in Normative Modeling

The core aim of normative modeling is to derive the normative range for a certain structural or functional brain measure. Therefore, we only need data from healthy participants to derive the model (although normative models can also be estimated from other populations). This property is advantageous given the excess of data availability for healthy populations compared to clinical populations. If successful, then any large deviation from this normative range is interpreted as an abnormality in the brain that can be studied concerning different mental disorders. These abnormalities can be quantified in the form of a probability by mapping each z-score *z* ∈ **z** in Eq. 1 to its corresponding probability using a modified cumulative distribution function of the normal distribution as follows:

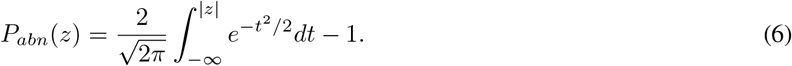

Here, we use this formulation because we consider, for example, samples in 1% and 99% percentiles equally abnormal. Other configurations can be adapted based on the definition of abnormality in a specific problem. The resulting probability is used as an abnormal probability index (*P_abn_*) for each sample. This index can be employed to detect anomalies in brain measures in an anomaly detection scenario (Kia and Marquand, 2018, 2019). This approach, in combination with normative modeling, provides an effective tool for data-driven biomarker discovery (see results in Sec. 3.4).

### 2.6 Experimental Material and Setup

In this section, we describe the experimental data and four experiments that are used for evaluating HBR in the normative modeling life-cycle.

#### 2.6.1 Datasets and Preprocessing

Table 1 lists the 16 neuroimaging datasets that are used in our experiments. For the ABCD dataset (Casey et al., 2018), we used data from the first imaging timepoint for subjects included in the *v*2.0.1 curated release. For the UK Biobank (UKBB) study (Miller et al., 2016), we used approximately 15000 subjects derived from the 2017 release. For the Human Connectome Project aging, development and early psychosis studies (HCPAG, HCPDV and HCPEP, respectively) we used data from the 1.0 data release. Further details surrounding the other datasets can be found in the relevant papers listed in Table 1. We have excluded participants with missing demographic (age/sex) information and those with poor quality imaging data. We excluded 1566 (4%) subjects due to low-quality images. Subjects were excluded if their scan-site median-centered absolute Euler number was higher than 25. The exclusion of outliers based on Euler numbers has been shown to be a reliable quality control strategy in large neuroimaging cohorts (Rosen et al., 2018; Sánchez et al., 2021). Median centering is necessary because the Euler number is scaled differently for different datasets. The threshold of 25 was determined empirically by manually examining the excluded scans. The final data consists of 37126 scans from 79 scanners that reasonably cover a wide range of human lifespan from 6 to 100 years old. Fig. 5 depicts the age span for each dataset. Note that the peak at approximately 10 years is driven by the ABCD dataset, where subjects are all nearly the same age. These properties make these data a perfect case-study for large-scale multi-site normative modeling of aging. The data also contain 1107 scans from participants diagnosed with a neurodevelopmental, psychiatric, or neurodegenerative disease, including attention deficit hyperactivity disorder (ADHD), schizophrenia (SZ), bipolar disorder (BD), major depressive disorder (MDD), early psychosis (EP), mild cognitive impairment (MCI), and (mild) dementia (DM).

**Table 1:** Demographics of multi-site experimental data. (*) The HCPDV and HCPAG datasets are collected by the same data acquisition centers. We consider this in computing the total number of scanners in data.

| Datasets | No. Scans | No. Patients | No. Scanners | Age Range | Sex M/F | FS Version |
| --- | --- | --- | --- | --- | --- | --- |
| ABCD (Casey et al., 2018) | 10732 | - | 29 | 9–11 | 52%/48% | 6.0 |
| CAMCAN (Taylor et al., 2017) | 647 | - | 1 | 18–88 | 49%/51% | 6.0 |
| CMI (Alexander et al., 2017) | 893 | - | 2 | 18–88 | 62%/38% | 6.0 |
| CNP (Poldrack et al., 2016) | 264 | 49(SZ),49(BD),41(ADHD) | 2 | 21–50 | 56%/44% | 6.0 |
| FCON (Biswal et al., 2010) | 1021 | 25(ADHD) | 18 | 8–85 | 43%/57% | 6.0 |
| HCP (Essen et al., 2012) | 1113 | - | 1 | 22–37 | 46%/54% | 5.3 |
| HCPAG (Bookheimer et al., 2019) | 677 | - | 5* | 36–100 | 43%/57% | 6.0 |
| HCPDV (Somerville et al., 2018) | 653 | - | 5* | 8–22 | 49%/51% | 6.0 |
| HCPEP (Seitz-Holland et al., 2021) | 180 | 123(EP) | 4 | 17–36 | 62%/38% | 6.0 |
| IXI (Imperial, 2021) | 557 | - | 1 | 20–86 | 44%/56% | 6.0 |
| NKI (Nooner et al., 2012) | 482 | - | 1 | 6–85 | 36%/64% | 6.0 |
| OASIS3 (LaMontagne et al., 2019) | 2044 | 271(DM),51(MCI) | 5 | 43–97 | 42%/58% | 5.3 |
| OPN (Stanford, 2021) | 612 | - | 6 | 8–58 | 45%/55% | 6.0 |
| PNC (Satterthwaite et al., 2016) | 1514 | - | 1 | 8–23 | 48%/52% | 6.0 |
| TOP (Skåtun et al., 2016) | 823 | 167(SZ),193(BD),31(MDD),107(others) | 1 | 17–69 | 53%/47% | 6.0 |
| UKBB (Miller et al., 2016) | 14914 | - | 2 | 44–80 | 48%/52% | 6.0 |
| **Total** | 37126 | 1107 | 79 | 6–100 | 49%/51% | - |

**Figure 5:**
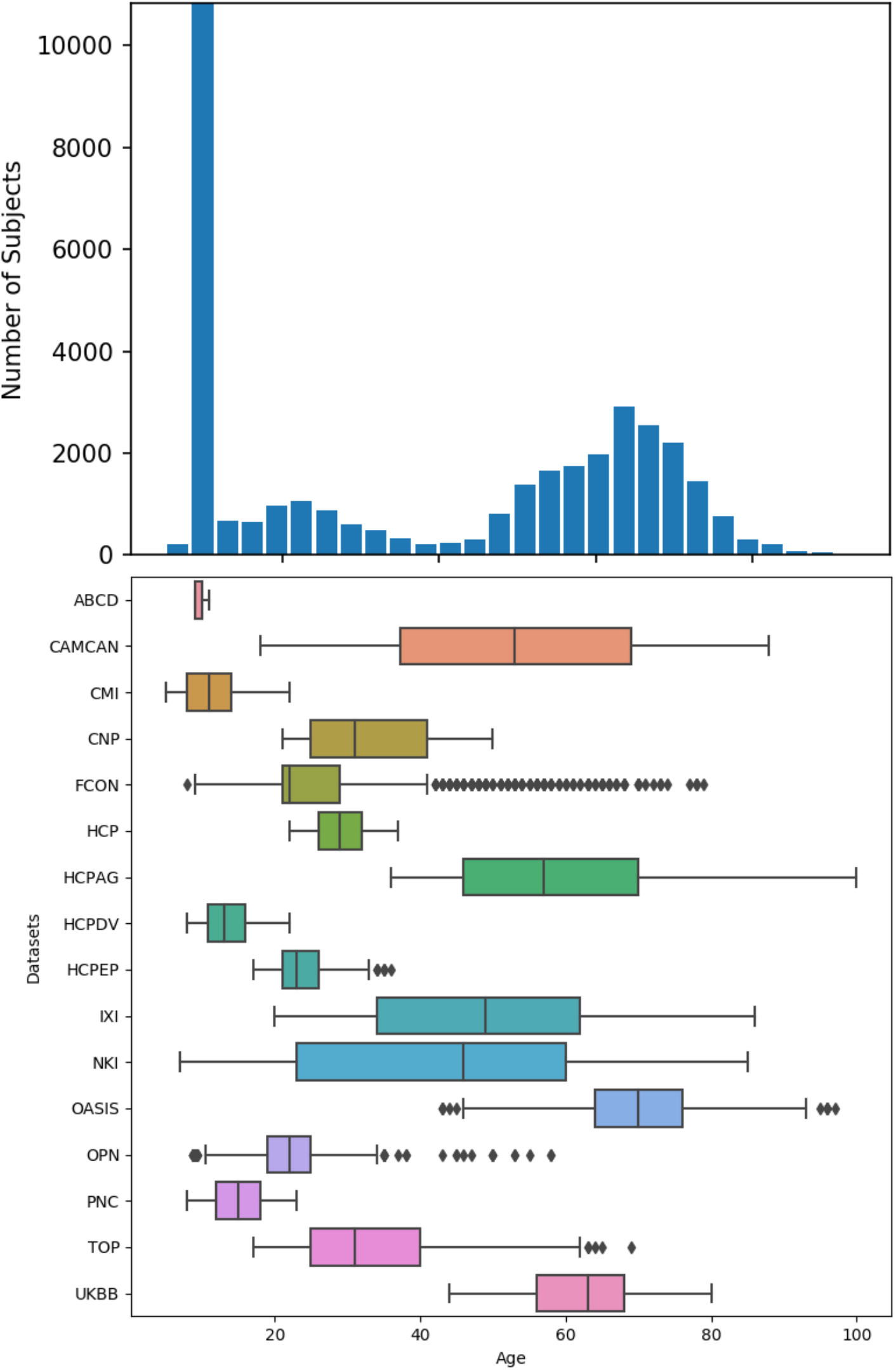
The age span of participants across 16 neuroimaging datasets. Our experimental data cover almost the full range of human life-span.

In our analyses, we use cortical thickness measures estimated by Freesurfer (Fischl, 2012) version 5.3 or 6.0 over 148 cortical regions in the Destrieux atlas (Destrieux et al., 2010). We have two motivations for this choice: i) the site-effect is very salient in the cortical thickness across data from different sites; ii) the fact that the effect of aging on thinning the gray matter is well-studied in the literature concerning different brain disorders. These features in data help us to better validate the method presented in this study. Fig. 6 shows the distribution of median cortical thickness with respect to age across participants and scanners. It clearly shows the presence of an overall effect of aging on cortical thinning and the site-effect in data. For example, the cortical thickness is on average higher in the UKBB dataset than in other datasets.

**Figure 6:**
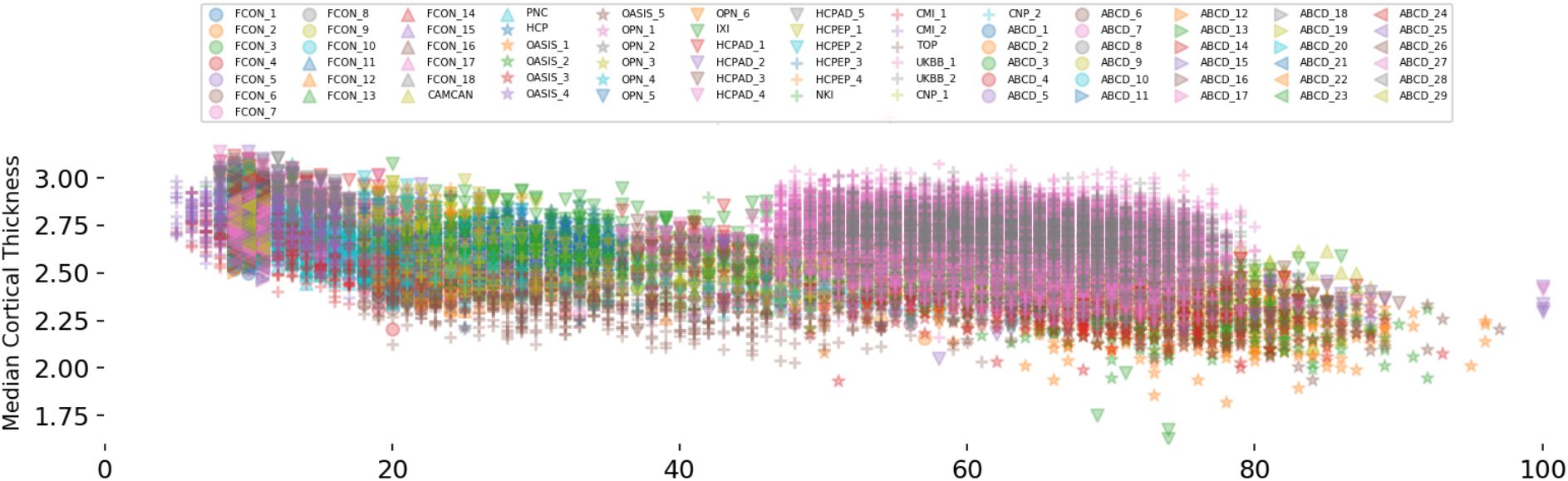
The distribution of median cortical thickness with respect to age across 79 scanners in 16 datasets. While an overall effect of aging on cortical thinning is present, however, it is highly contaminated with site-effect. The data in some datasets (*e.g.*, UKBB) show relatively higher cortical thickness compared to the others.

#### 2.6.2 Experiments

To demonstrate the effectiveness of HBR in completing the life-cycle of large-scale normative modeling, we set up four experimental settings for predicting the cortical thickness across 148 cortical regions: 1) multi-site data regression, 2) model extension, 3) model adaptation, and 4) anomaly detection. In all experimental configurations, we use only age as a covariate except for the fixed-effect site modeling in which, by definition, the one-hot encoding of scanner ids are also included in the covariates. We use sex as a group-effect in all estimated models. In the HBR case, the scanner is also included as a group-effect. All experiments and evaluations are repeated ten times with different random healthy participants in the training and test phases.

In the multi-site data regression setting, the goal is to compare the performance of HBR with naive pooling, fixed-effect pooling, pooling after data harmonization, and no-pooling models in deriving the normative range of cortical thicknesses across 148 brain areas. Here, we assume the data from all scanners are available when estimating the normative model, *i.e.*, a centralized data architecture. In each experimental run, 80% of healthy samples are randomly selected to train the regression model. The remaining 20% are used for the evaluation. We modeled the effect of age on the response variable in three ways, 1) as a linear effect, 2) as a non-linear effect using a cubic polynomial, 3) as a non-linear effect using a cubic B-spline basis set expansion with 5 evenly spaced knots. Given the characteristics of experimental data, we use a site-specific homoscedastic form for the variance. We emphasize that the proposed framework is capable of modeling heteroscedasticity. Here, using a heteroscedastic model for the variance did not provide any advantage at the cost of higher model complexity (see the supplementary for the comparison). Three metrics are used to evaluate the quality of fits, i) Pearson’s correlation coefficient (RHO) between observed and predicted brain measures; ii) standardized mean squared error (SMSE), and iii) mean standardized log-loss (MSLL). In the latter two cases, the lower values for the metrics represent the higher quality of the fitted function. While correlation and SMSE evaluate only the predicted mean, MSLL also accounts for the quality of estimated variance which plays an important role in deriving deviations from the norm (see Eq. 1).

In the model extension experiment, the goal is to compare the performance of the HBR models trained on the centralized and decentralized data. While in the former, we use the same configuration in the multi-site regression experiment, in the latter case, we assume that we have only access to one dataset at each time-step and we estimate the model parameters sequentially, *i.e.*, adding one dataset at a time until it covers all datasets. We generated 5 samples for each age value (in the range of 10 to 90 years old) and each gender in the data generation process for each dataset (80 × 2 × 5 = 800 samples). The same evaluation metrics are used to compare these two different settings.

In the model adaptation setting, we demonstrate an application of HBR in a more realistic clinical scenario when the aim is to adapt the parameters of a reference normative model to private clinical data at local hospitals. To do so, we first use a linear homoscedastic model to estimate the parameters of the reference normative model on datasets with only healthy participants (ABCD, CAMCAN, CMI, FCON, HCP, HCPAG, HCPDV, IXI, NKI OPN, PNC, and UKBB). Then, in each run 50% of random healthy participants in clinical datasets, including CNP, HCPEP, OASIS3, and TOP are used to recalibrate the parameters of the reference model. The rest of the healthy participants and patients are used as test samples. It is important to emphasize that other methods including harmonization and complete pooling do not apply to this setting because they do not support model adaptation to new datasets. We compare the HBR with no-pooling in which separate models are trained for each clinical dataset.

In the anomaly detection experiment, we aim to exemplify a possible application of the full cycle of normative modeling (*i.e.*, developing a reference normative model on a large healthy population and model adaptation to clinical data) in data-driven biomarker discovery. Here, we use the resulting z-scores in the model adaptation experiment in the anomaly detection scenario described in Sec. 2.5. The abnormal probability indices for each individual across 148 cortical regions are computed. Then, the region-wise areas under the ROC curves (AUCs) are derived to evaluate the predictive power of deviations for each diagnostic label. We employed a conservative approach to testing for statistical significance, where we performed permutation tests with 1000 repetitions and used false discovery rate (FDR) correction (Benjamini and Hochberg, 1995) to correct for multiple comparisons across 148 regions. To ensure the stability of results, only significant areas that pass the FDR correction in 9 or more out of 10 full experimental runs are reported. We refer to this as ‘significant and stable’.

#### 2.6.3 Implementations and Model Settings

The HBR model is implemented in Python using the PyMC3 package (Salvatier et al., 2016). A No-U-Turn sampler (NUTS) (Hoffman and Gelman, 2014) is used for inferring the posterior distributions of parameters and hyperparameters. Normal and log-normal distributions are respectively used as hyperpriors for the mean and standard deviation of parameters of *f_μ_* (see Fig. 3). The distribution of the standard deviation of the homoscedastic noise in logarithmic space is set to a normal distribution with 0 mean and standard deviation of 2.5. Non-centered parameterizations are used to simplify posterior geometries and increase the performance of the sampler (Betancourt and Girolami, 2015). All implementations are available online within the PCNToolkit *v*0.18 package at https://github.com/amarquand/PCNtoolkit. The high-performance computing techniques are employed in our implementations to parallelize the computations across computational nodes on a computer cluster.

For harmonizing data using ComBat, we use a Python implementation available at https://github.com/Warvito/neurocombat_sklearn. This implementation has the possibility to learn the ComBat parameters on the training data and apply them to the test data that is and essential feature for out-of-sample evaluations in our experiments. Age and sex are used in the design matrix (**X** in Eq. 3) when applying the ComBat for data harmonization to ensure that their variability is preserved in data (see the distribution of data after harmonization in the supplementary material).

## 3 Results

In this section, we present the results of four experiments in Sec. 2.6.2.

### 3.1 HBR, Suitable Flexibility for Big Multi-site Data

Fig. 7 summarizes the empirical densities over three evaluative metrics (across 148 cortical areas) in the multi-site data regression scenario. Each column compares an evaluation metric across five modeling approaches (naive pooling (NV), fixed-effect pooling (FE), pooling after ComBat harmonization (CMB), HBR, and no-pooling); and three different model parametrization for the mean effect (linear, polynomial, and B-spline). In all cases, the HBR and fixed-effect modeling show equivalently better regression performance compared to other approaches. These two models both account for site-effect in data (unlike naive pooling), benefit from the richness of big data (unlike no-pooling), and have enough model flexibility to find the best fit to data (unlike naive pooling and pooling after harmonization). Even though they use different strategies to provide this flexibility; HBR by accounting for the difference between the distributions of signal and noise across multiple sites rather than ignoring or removing it, and the fixed-effect pooling by increasing the degree-of-freedom of the model via additional covariates (80 versus 1 for other models). However, this increased flexibility may result in an inferior performance when applied to small sample-size data. In addition, in fixed-effect modeling, as discussed in Sec. 2.3.3, using batch-effects as covariates in the fixed-effect pooling method may result in regressing out informative but *a-priori* unknown variance from data.

**Figure 7:**
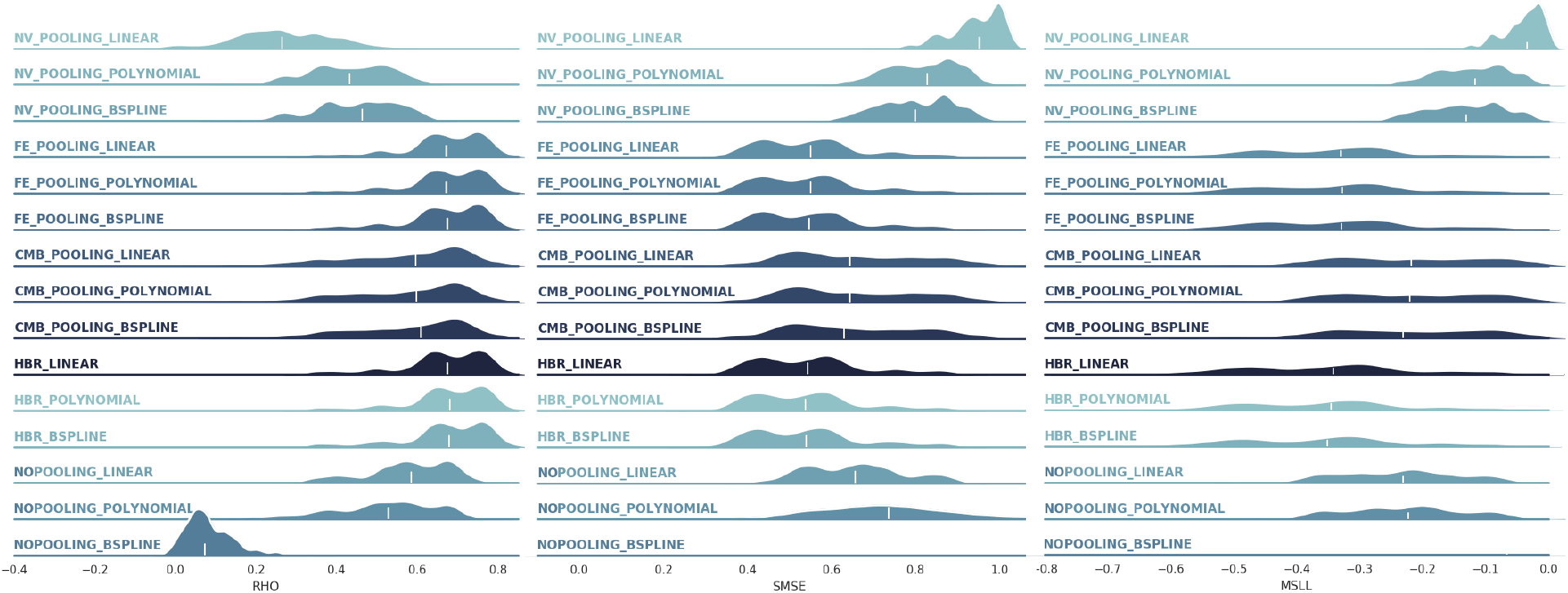
The distributions of correlation, SMSE, and MSLL across 148 cortical areas in the multi-site data regression. The white lines highlight the medians of distributions. Abbreviations: NV=naive, FE=fixed effects, CMB=ComBat, HBR=hierarchical Bayesian regression. The HBR and fixed-effect modeling show equivalently better regression performance compared to other approaches.

On the other hand, on these experimental data, using more complex non-linear parametrizations has shown a negligible positive effect on the performances and linear models still provide competitive results. The very poor performance of the B-spline model in the no-pooling model (the SMSE and MSLL are out of range of plots) is the consequence of over parametrization on small sample size data. In short, our results show that choosing the right model flexibility on bigger data always results in more favorable regression performances. Therefore, taking an appropriate strategy for handling the site-effect in data can play a vital role in finding a better fit to data resulting in a better estimation of the normative range in normative modeling. Our experimental results confirm that HBR affords the proper model flexibility for modeling big multi-site data.

The distribution of measured metrics across 148 brain regions (thus 148 models) is multi-modal and wide in some cases, especially for MSLL measures. These diverse results across brain regions can be explained from data and model perspectives. From the data perspective, some brain regions might have a lower or higher relationship with covariates of interest (*e.g.*, age). When there is no relationship between the two sides even the most complex models will fail if fairly evaluated. From the modeling perspective, in some brain regions, the model may not be able to explain the relationship between covariates and target brain measures. For example, when using linear models for modeling non-linear relationships. In this experiment, since all linear, polynomial, and B-spline models show similar performance, we conclude that the low performances (in MSLL, SMSE, and RHO) in some brain regions are due to the small effect of aging on the cortical thickness.

We conducted an extra experiment to evaluate the performance of different techniques in removing site-effects in resulting z-statistics. In this experiment, the z-statistics are computed on the test sets across different modeling approaches. Then they are used as input to a linear support vector machine (SVM) classifier (C=1) to classify the datasets in a one-vs-one scenario. The balanced-accuracies computed over 5-fold stratified cross-validation are reported in Fig. 8. The meaningful difference between classifier performances of naive pooling (that ignores the site-effects) and the other methods (that use different strategies to exclude site-effects) demonstrates 1) the importance of correcting for site-effects; 2) the effectiveness of these different strategies to remove a majority of site variation in data (drop in the average accuracy from 0.90 to ∼ 0.53).

**Figure 8:**
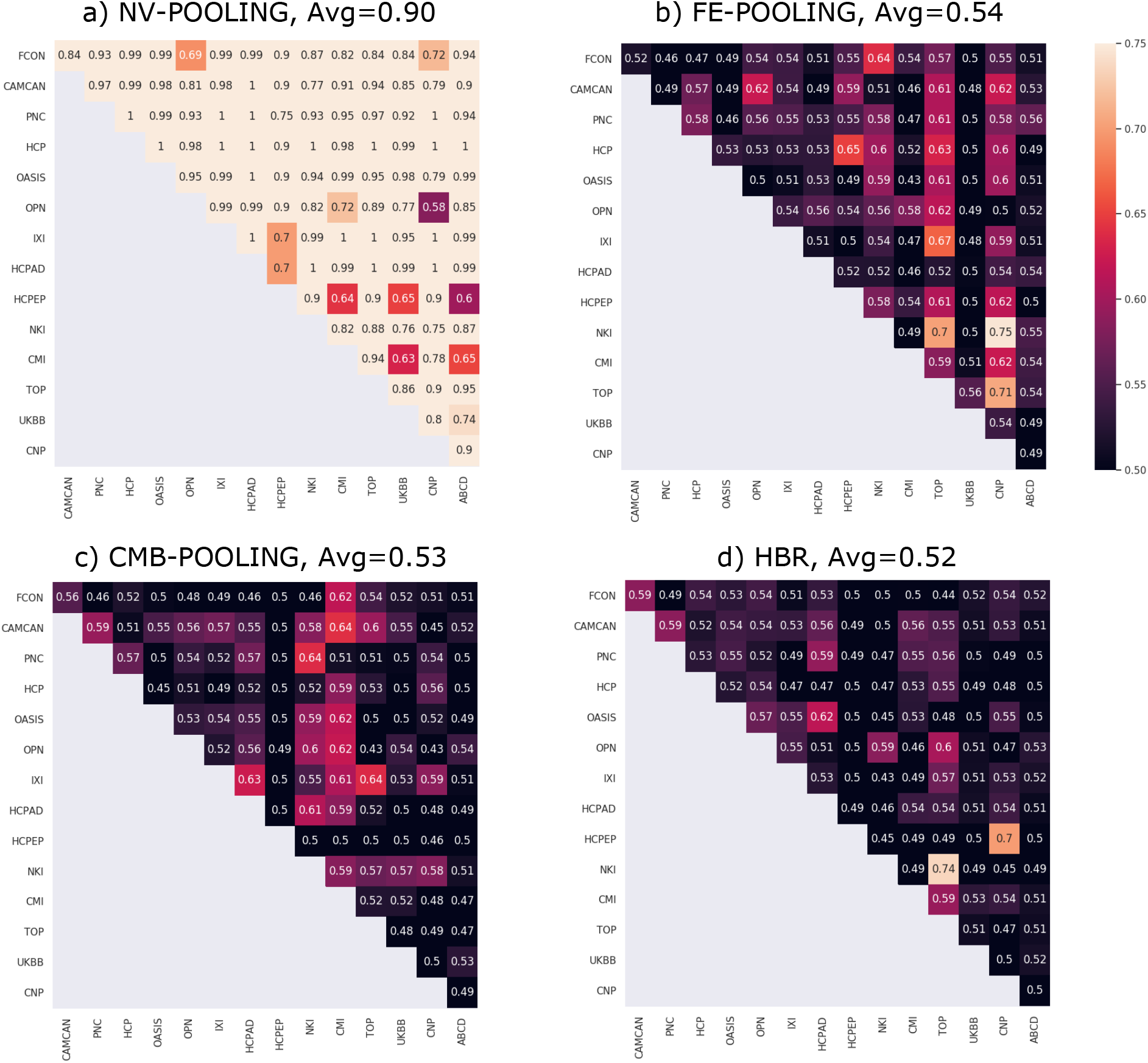
Balanced-accuracies in classifying z-statistics across different datasets in a one-vs-one scenario. The z-statistics are computed using different modeling approaches including naive pooling (NV-POOLING), fixed-effect pooing (FE-POOLING), pooling after ComBat harmonization (CMB-POOLING) and HBR. The results show that the site-effects are to high degree not present in the z-statistics in FE-POOLING, CMB-POOLING, and HBR.

### 3.2 HBR, Distributed Modeling on Distributed Data

Fig. 9 compares the evaluation metrics for the linear HBR model with homoscedastic noise when trained on the centralized and decentralized multi-site data, adding one site at a time. The extended model shows very close RHO, SMSE, and MSLL compared to the model trained on full data in one run (*R*^2^ = 0.98, 0.97, 0.95, respectively). These results show the success of the proposed model extension strategy in estimating the mean prediction. However, the MSLL measure shows a slight but negligible decline in some regions; revealing the lower performance of the extended model in capturing the actual variance in some brain areas. Generating more samples in the data generation process might improve the model quality from this respect at the higher computational costs in time and memory. These promising results confirm the possibility of estimating multi-site normative models on distributed data across multiple data centers. This can significantly reduce the need for sensitive clinical data sharing. Furthermore, it reduces the data transfer, maintenance, and storage costs for storing several copies of the same data across several centers in centralized model development.

**Figure 9:**
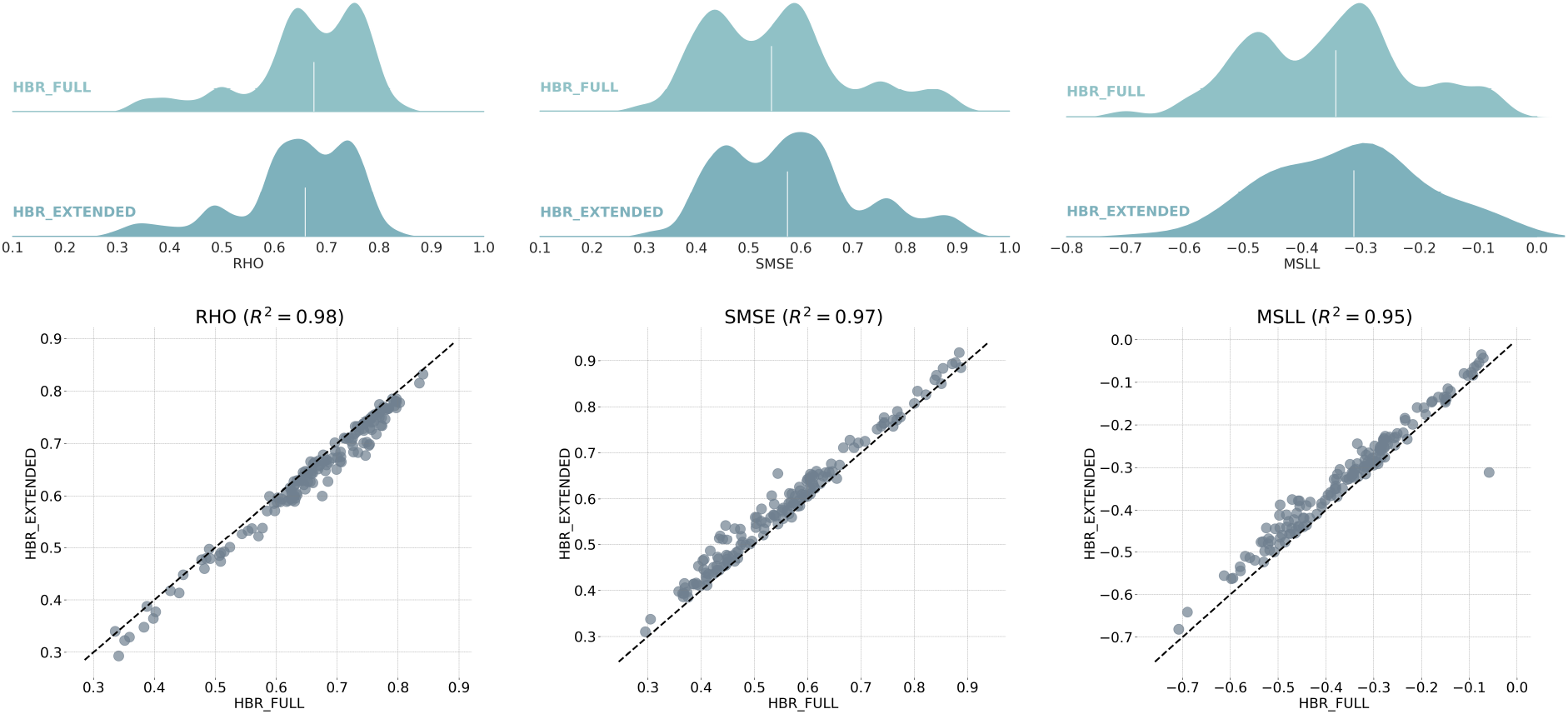
Comparison between the regression performance of HBR when trained on centralized data (HBR-FULL) versus decentralized model development using model extension strategy (HBR-EXTENDED). The ridge plots show distributions of correlation, SMSE, and MSLL across 148 cortical areas. As depicted in the scatter plots, HBR-EXTENDED models show very similar RHO (*R*^2^ = 0.98), SMSE (*R*^2^ = 0.97), and MSLL (*R*^2^ = 0.95) compared to HBR-FULL models trained on full data in one run.

### 3.3 Prior Information Matters

Fig. 10 compares the regression performance of the no-pooling approach with the adapted HBR model. While in the first case, we separately model the data from each presumably clinical center, in the second case, we try to benefit from transferring the knowledge from the reference normative model to local models. In all three evaluative metrics, the adapted HBR model shows a better regression performance compared to no-pooling. These results confirm the value of prior information learned by the reference model on big data in estimating more accurate normative models.

**Figure 10:**
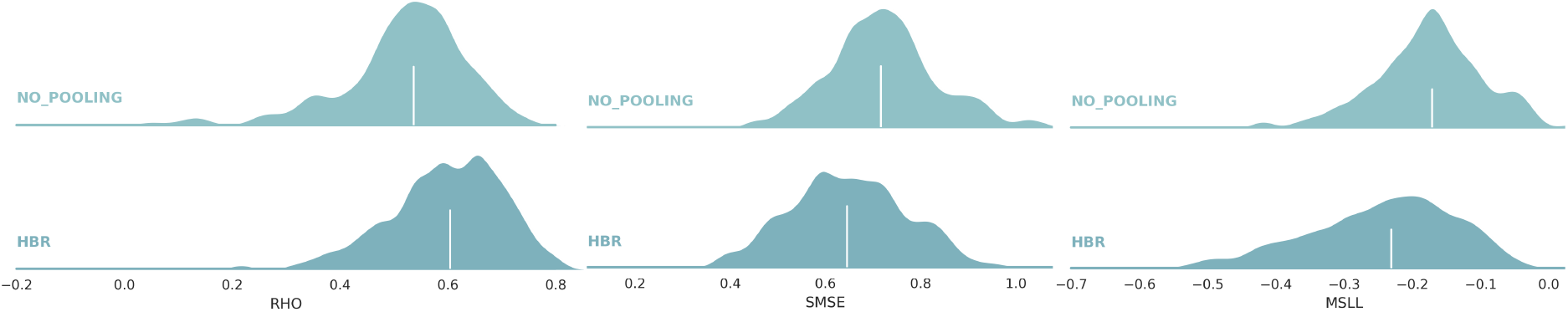
Comparing the regression performance of the adapted HBR model versus the no-pooling strategy. The ridge plots show distributions of correlation, SMSE, and MSLL across 148 cortical areas. The adapted HBR model shows a better regression performance compared to no-pooling.

### 3.4 The Deviations are Distinctive

In Fig. 11, we depict significant and stable AUCs across brain regions for different complex brain disorders and diseases. Here, the procedure explained in Sec. 2.5 is used to derive the abnormal probability index for each sample and each region. Only significant and stable areas (see Sec. 2.6.2 for more detail on significance and stability criteria) are reported. Only the results for dementia, schizophrenia and early psychosis could pass our rigorous stability test.

**Figure 11:**
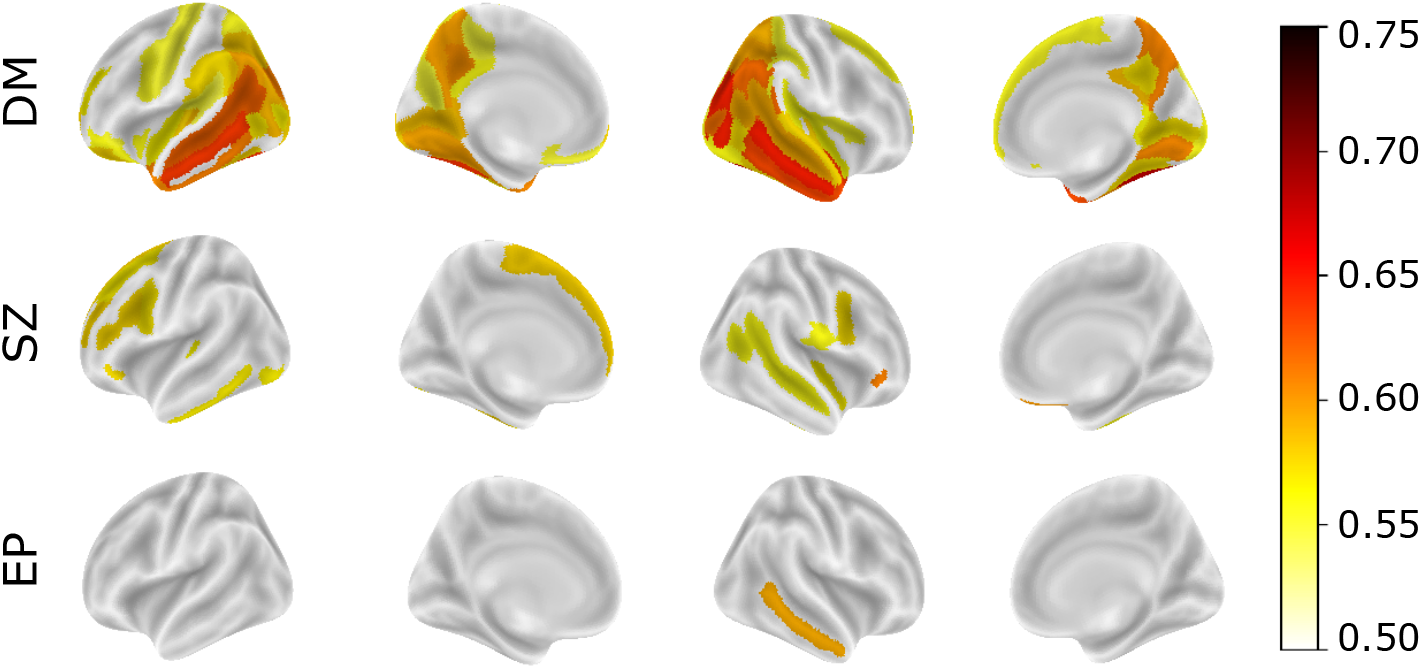
Significant and stable AUCs across brain regions for detecting healthy participants from patients in the anomaly detection scenario. In dementia (DM), the best performances are observed in the occipital and temporal lobes including bilateral occipitotemporal (fusiform) gyrus, right middle temporal gyrus, right superior/transverse occipital sulcus, right middle occipital gyrus, and left middle temporal gyrus. In schizophrenia (SZ), the distinctive areas are in the frontal lobe including the right orbital inferior frontal gyrus, right medial orbital sulcus, left superior frontal gyrus, left middle frontal sulcus, left anterior transverse collateral sulcus, and left inferior frontal sulcus. In early psychosis (EP), only the right middle temporal gyrus shows significant and stable AUC.

In dementia cases, the best performances are observed in the occipital and temporal lobes including bilateral occipitotemporal (fusiform) gyrus (AUC=0.68 and 0.64), right middle temporal gyrus (AUC=0.65), right superior/transverse occipital sulcus (AUC=0.65), right middle occipital gyrus (AUC=0.65), and left middle temporal gyrus (AUC=0.64). Fig. 12 shows that patients with dementia manifest relatively stronger deviations in the respective brain regions. The proposed approach is capable of detecting brain regions that are repeatedly reported in the literature and have been linked to dementia (Yang et al., 2019a; Soheili-Nezhad et al., 2020; Machulda et al., 2020; Habes et al., 2021). Note that the patients with dementia were derived from the OASIS3 dataset, which contains only mild cases. Therefore, the accuracies reported are not directly comparable with studies derived from patients with more advanced forms of dementia.

**Figure 12:**
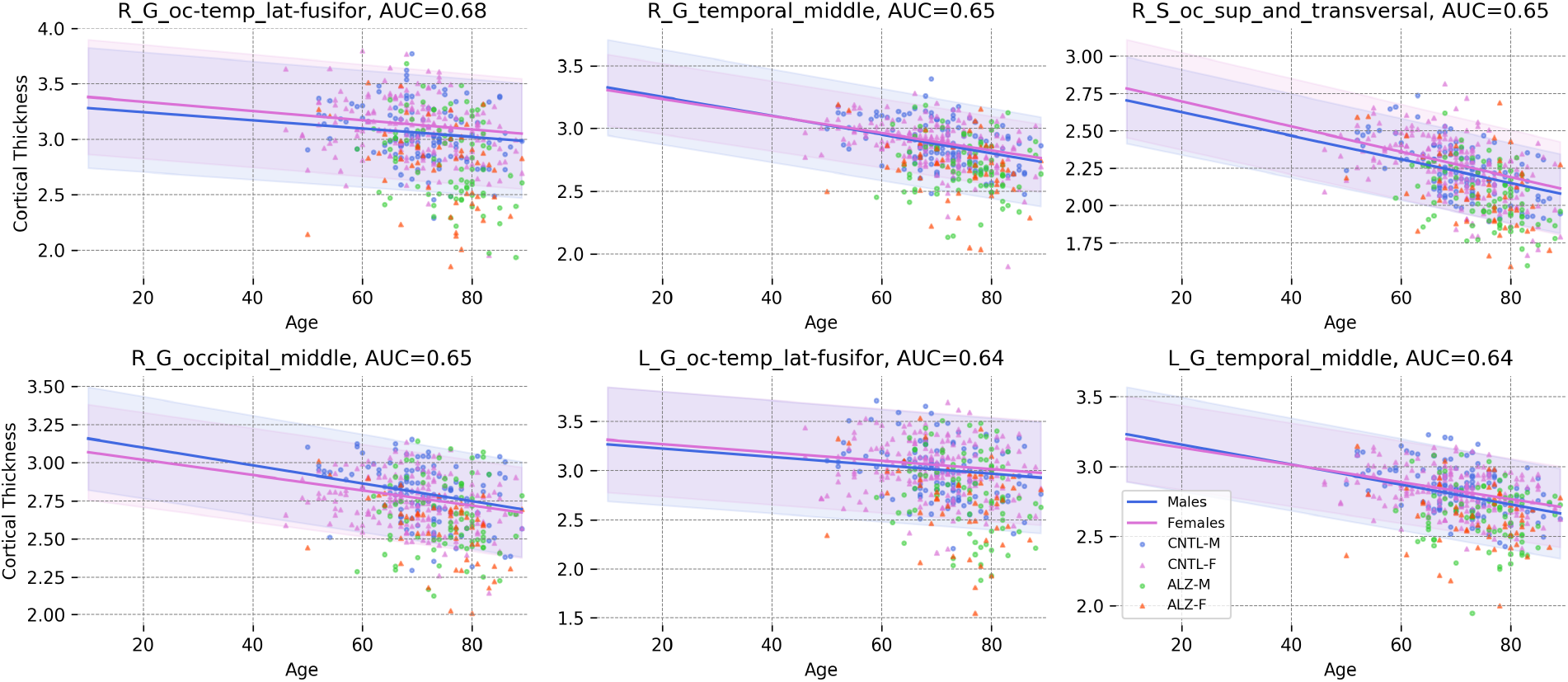
The norm and the 95% normative range for males and females in the 6 most distinctive cortical regions in dementia. Patients show lower cortical thickness than the norm of the population (Abbreviations: CNTL-F=healthy female, CNTL-M=healthy male, DM-F=female patient, DM-M=male patient).

In patients with schizophrenia, our results show the concentration of distinctive areas in the frontal lobe including the right orbital inferior frontal gyrus (AUC=0.61), right medial orbital sulcus (AUC=0.60), left superior frontal gyrus (AUC=0.58), left middle frontal sulcus (AUC=0.58), left anterior transverse collateral sulcus (AUC=0.58), and left inferior frontal sulcus (AUC=0.58). These results are compatible with previous studies reporting cortical thinning in the frontal lobe in patients with schizophrenia (Rimol et al., 2010, 2012; van Erp et al., 2018). Fig. 13 shows how the cortical thicknesses of patients in these areas are distributed around the normative range. In early psychosis, only the right middle temporal gyrus (AUC=0.59) shows significant and stable results. Cortical thinning in temporal regions has been reported in earlier studies on EP patients (Sumich et al., 2002; Makowski et al., 2016; Roalf et al., 2016).

**Figure 13:**
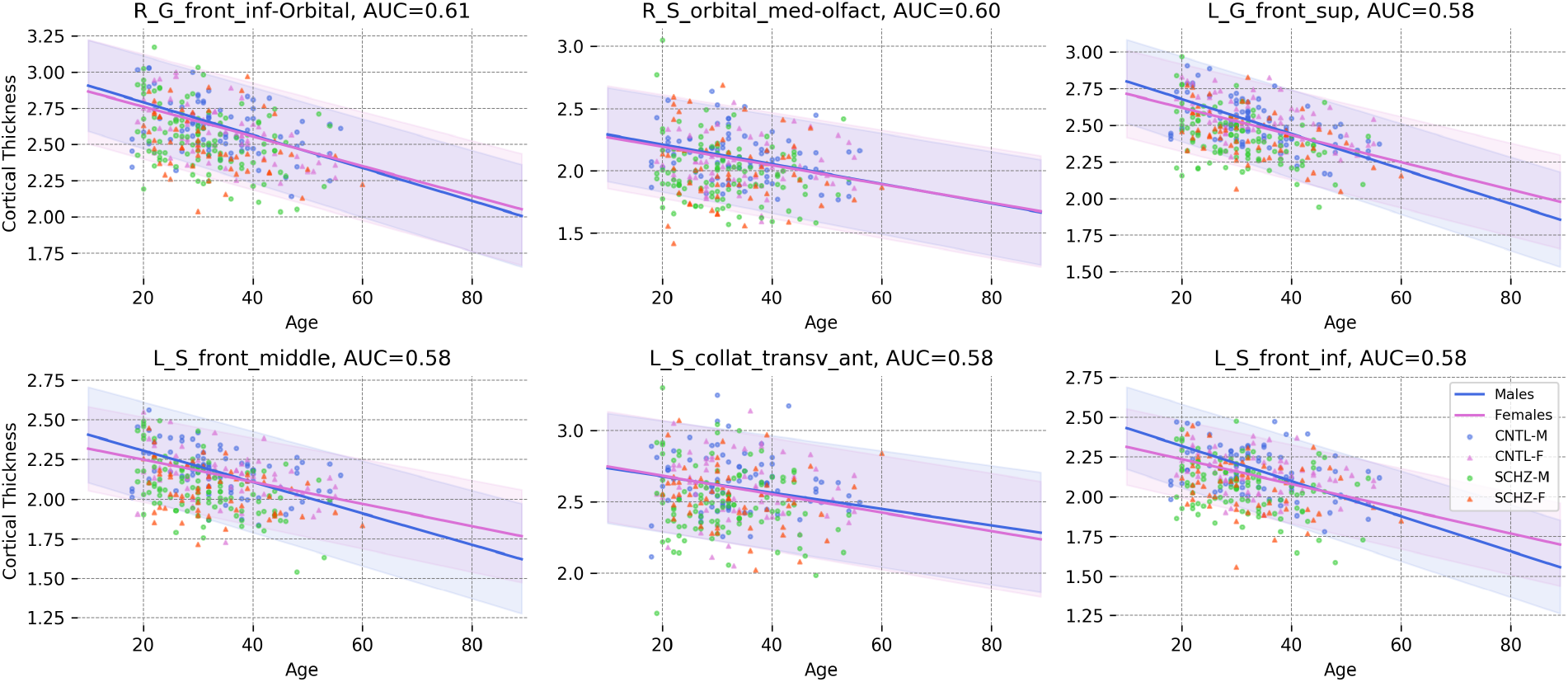
The norm and the 95% normative range for males and females in the 6 most distinctive cortical regions in schizophrenia. Patients show lower cortical thickness than the norm of the population (Abbreviations: CNTL-F=healthy female, CNTL-M=healthy male, SZ-F=female patient, SZ-M=male patient).

Even though these performances are lower than the state-of-the-art in classifying dementia and schizophrenia patients from healthy participants, it is important to consider the fact that our anomaly detection method is, in contrast, an unsupervised approach; in the sense that the model does not see any patient data during the training phase (*i.e.*, in deriving the normative range).

Since patterns of sub- and supra-normal deviations can be extracted on an individual basis (Wolfers et al., 2018, 2020, 2021), our approach can be used as a tool for precision psychiatry by decoding the heterogeneity of complex brain disorders at the level of an individual patient (Foulkes and Blakemore, 2018; Marquand et al., 2019).

#### 3.4.1 Few-Shot Learning on Small Data

Another appealing experimental observation in the model adaptation setting is the potential of the HBR model in learning reasonable normative ranges on tiny datasets. Fig. 14 shows the normative range for the right middle temporal gyrus (the most distinctive region for early psychosis) across four different HCPEP acquisition sites. Only healthy subjects in the training set are depicted in the plots. In all sites, only a few training subjects are available for estimating the normative model. However, the HBR model can still find a reasonable estimation of normative ranges even for the extreme cases in the second and the third sites in which, respectively, 0 and 1 training samples are available.

There is a key feature in the HBR framework that contributes to this performance: the informative prior that the adapted model inherits from the reference normative model. This informative prior is already learned from thousands of data points and acts as a high-level regularizer that prevents the model parameters from overfitting to misleading data. The example is the best linear fit for females (dashed purple line in the top left plot in Fig. 14). Without having prior knowledge about the underlying effect of aging on cortical thickness, the best linear estimate on the training data shows ascending trend for cortical thickness with aging. Another example is tiny training data with 1 or even 0 samples (like the females in the second and third sites). In these cases, estimating the parameters of the linear model is impossible, thus the prior knowledge about the problem plays a decisive role in finding reasonable estimations. We emphasize that this is not only a theoretical problem because, in practice, multi-site clinical datasets often have sites with few samples. These results demonstrate the capabilities of the HBR model for few-shot learning (Wang et al., 2020) when adapting the reference model to very small local datasets. This feature can play even a more crucial role when adapting more complex normative models (for example, when *f_μ_* and *f_σ_* are parametrized on a neural network) on small data at local clinical centers.

**Figure 14:**
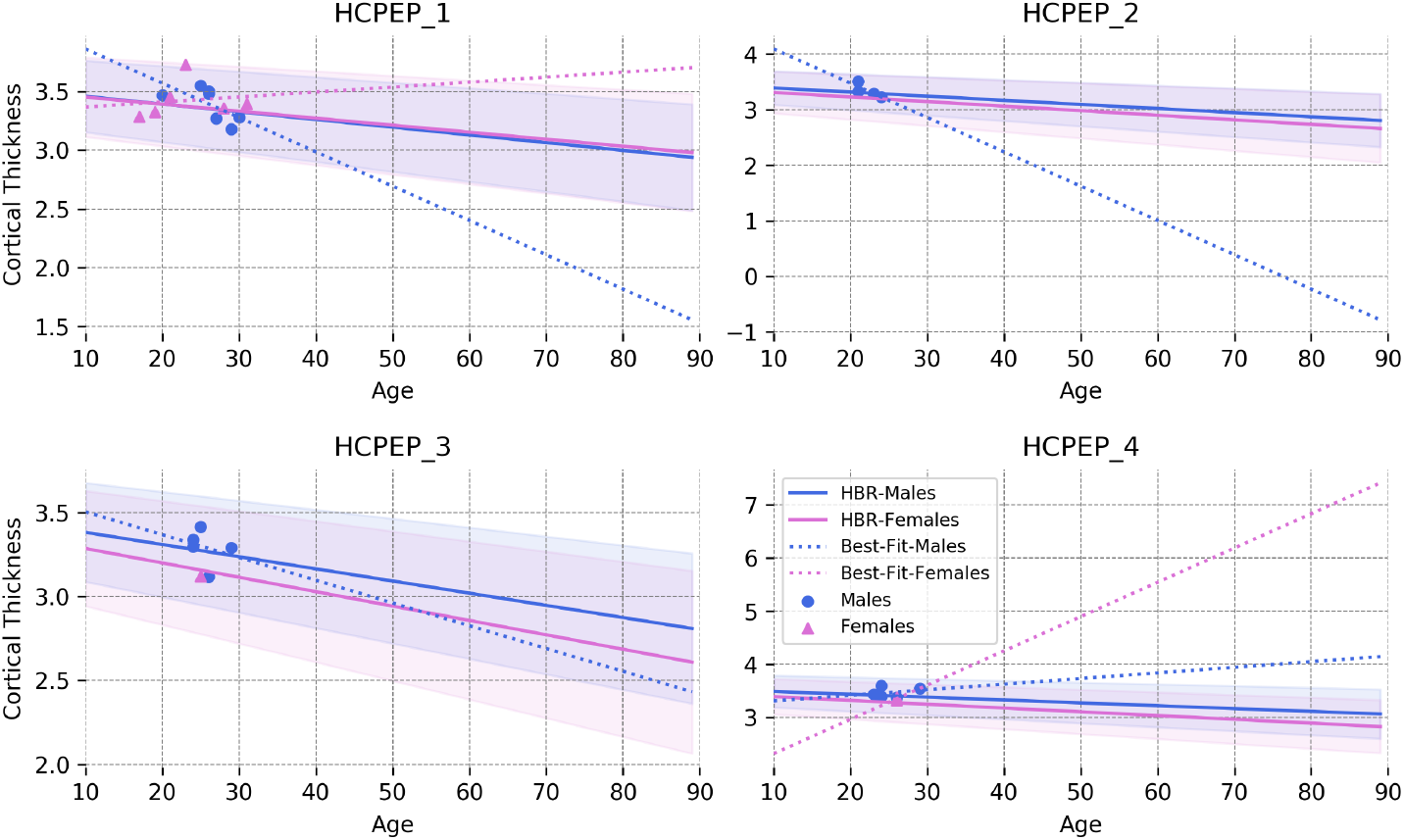
Normative ranges for the right middle temporal gyrus estimated by the adapted HBR model across four sites in the HCPEP dataset. The data points in the plots show the healthy subjects in the training set. The dashed lines show the best linear fit to the points for each sex. Benefiting from informative priors, the adapted HBR model provides a reasonable estimate of the normative range even on tiny training data.

## 4 Discussion

Our positive experimental results demonstrate the success of hierarchical Bayesian modeling in fulfilling the technical demands for closing the life-cycle of normative modeling. In the following, we will discuss the methodological significance and the clinical relevance of our contributions. We further pinpoint the limitations of the proposed method and envisage possible directions for future enhancements.

### 4.1 The Methodological Significance

Our HBR approach provides practical solutions to several key problems necessary to close the loop of normative modeling on realistic population-scale clinical datasets. Our main contributions are: i) accurately estimating centiles of variation whilst properly accounting for site variation with ii) manageable computational scaling to massive neuroimaging datasets; iii) a federated learning life-cycle that allows models to be updated as new datasets become available, without requiring access to the primary data and iv) enabling the transfer of information from population-level datasets to small clinical datasets.

We have designed our approach from the ground up with real-world clinical datasets in mind. The federated and distributed nature of our architecture is very important because it allows us to use large publicly available datasets for charting variation across the population to extract maximal value from clinical datasets that are often small and acquired on specific scanners. We consider model portability to be important for clinical applications. It is not feasible to transfer hundreds of thousands of scans to make predictions at a clinical site and –conversely– many clinical datasets are still small and can also be difficult to transfer (*e.g.*, if subjects contributing data did not provide the necessary consent).

In this work, we have considered only models that are linear in the parameters. More specifically, for most of the experiments, we parameterized the HBR method as linear, allowing the estimation of a different slope for each scan site. While non-linear effects are seen in some neuroimaging derived measures (Ziegler et al., 2012). Our results suggest that the linear model is sufficient for cortical thickness, and non-linear basis expansions did not explain more variance. However, for other neuroimaging-derived measures non-linear or heteroscedastic models may be more appropriate. We emphasize that our approach is fully modular, and such extensions can be easily integrated by adjusting the parametrization (see for example (Bayer et al., 2021)).

Taken together, HBR provides a principled, complete, and unified solution for modeling inter-subject variation in big data cohorts.

### 4.2 The Clinical Relevance

In our clinical application, we show that patients deviate significantly more from the estimated norm than healthy individuals (Fig. 12 and 13) with a regional distribution of abnormalities that is largely consistent with the known pathology of each disorder. This is in line with earlier publications (Wolfers et al., 2018; Zabihi et al., 2019; Wolfers et al., 2020; Zabihi et al., 2020; Wolfers et al., 2021), which show that while we observe differences between groups of patients and controls, those differences are not perfect to the extent of complete group separation (Marquand et al., 2019). This has been linked to the heterogeneous nature of these illnesses, which generally show a unique pattern of sub- and supra-normal deviations in individuals even when diagnosed with the same illness.

The ability of our approach to estimating normative models without the requirement to share sensitive data across different imaging and clinical centers can not be overemphasized in value as it allows us to map differences between individuals with a complex illness on a previously unprecedented scale. This is important as complex brain disorders are believed to have a unique manifestation across individuals (Foulkes and Blakemore, 2018). Therefore, it is necessary to map those differences in large samples. Here, we provide a framework and tool that will allow us to extend this work in a principled fashion towards multi-center imaging studies such as the ENIGMA consortium (Thompson et al., 2014). While the present paper has a technical focus, we can already show that our method is capable to detect significant deviations from a normative process in individuals with a complex illness. Our contributions pave the way toward incorporating biological measures into the diagnosis and treatment of mental disorders to hopefully find the right treatment at the right time for the right patient.

### 4.3 HBR versus Data Harmonization

In this study, we proposed an application of hierarchical Bayesian regression (HBR) for specific usage in federated multisite normative modeling. In a normative modeling setting, we presented experimental evidence for the effectiveness of our method in deriving more accurate normative ranges and mitigating site-effects in resulting statistics. We showed how the HBR can be used as an alternative to data harmonization and fixed-effect modeling by resolving their theoretical and practical limitations in multi-site normative modeling on decentralized data. Nevertheless, we must emphasize that we do not consider HBR to be a data harmonization method, *per se*. Therefore, if the aim is merely data harmonization for other purposes than normative modeling then the HBR is not an appropriate choice because, whilst the z-statistics are cleared of site-effects, site-related variance is still present in the HBR predictions (*f_μ_*).

One of the important differences with respect to most harmonization techniques is that HBR enables estimating sitespecific mean effects (*f_μ_*) and variations 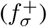 which are used in the normative modeling context to derive site-agnostic z-statistics. In contrast to most harmonization techniques, which often pool estimates over voxels or regions of interest, HBR pools over sites. This allows each site to have a different relationship with the covariates (*e.g.*, different slopes or variances, as illustrated in supplementary Fig. S2 and S3). This provides several advantages: first, it preserves differences across the range of the covariates (*e.g.*, increasing variance with age across the lifespan in scenarios where age is correlated with site-effect), rather than forcing each site to have the same average variance. Second, it allows transfer learning to new sites, where the parameters are adjusted according to the characteristics of the new site and regularised by the informative prior distribution learned across the original sites, providing increased flexibility over (for example) harmonizing the data by applying the parameters learned on one set of data to a new dataset. This procedure is similar in spirit to meta-analysis as the second level parameters of the model (*θ_μ_* and *θ_σ_*) and z-statistics are estimated for each site separately (but not independently). On the other hand, it is also similar to mega-analysis because the first level parameters (the parameters of the prior including 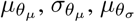, and 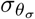) are estimated jointly across sites (See Fig. 3).

In contrast, whilst harmonization provides the possibility to merge the data across different centers and perform the analysis on pooled data, this process might be harmful in the normative modeling context (see Sec. 2.3.2 and our simulation study in the supplementary material) in which we are interested in the exploratory analysis of the variation in data. With this in mind, we do not claim HBR is a complete alternative to harmonization, and we recommend users choose the optimal approach according to their specific analytical goals.

### 4.4 Limitations and Future Directions

The current implementation of HBR employs a Gaussian likelihood function, hence, assumes a Gaussian distribution for residuals. If this is not the case, the estimated centiles and z-scores might not be well-calibrated. Although this is usually not a big problem, distributions of some phenotypes are known to be skewed or bounded, and in these situations, this method would not provide accurate results. However, the presented HBR method is fully capable of accommodating non-Gaussian variability in data. This possibility can be easily implemented by changing the likelihood and parametrize it over location, scale, and shape parameters (instead of mean and variance). Because we use a sampling approach for the inference, our method can estimate complex non-trivial posterior distribution with no closed-form analytical solutions. We are currently working on finalizing this extension. Another possible direction to solve this problem is to use likelihood warping (Fraza et al., 2021), in which the data with arbitrary distribution is first warped into a Gaussian distribution and ordinary methods (with normality assumption) can be applied to derive the centiles of variation. Then these centiles are transferred back to the original distribution using a reverse operation.

Another limitation is that models are estimated separately for each brain region without accounting for correlations between brain regions. While this removes nearly all univariate site variation, our results in Fig. 8 show that in all methods (including fixed-effect pooling, pooling after ComBat harmonization, and HBR), still, a few considerably above the chance-level performances are present in the tables. This is mainly because, in all benchmarked models, the harmonization and modeling are performed separately for each brain region without accounting for correlations between brain regions. Hence, only univariate site-effects are removed, and still, some multivariate site-effect might be present in data. Therefore, machine learning classifiers could still learn this residual information from the data (Dinga et al., 2020). One possible remedy for this problem is to harmonize the covariance structure of multivariate data (Chen et al., 2020). Another option is to remove the batch-effects from machine learning predictions (Dinga et al., 2020). Of course, the presented anomaly detection method is immune to this deficit because it is separately applied to each cortical region. A conceptually straightforward extension to our model is to model correlations between brain regions that are related to site-effects in data. This extension can be straightforwardly integrated into the present model, for instance, using Wishart priors for the covariance between brain regions.

The quality of scans is a crucial factor in the success of developing a reference normative model. Low-quality noisy scans can impair the inference and result in inaccurate estimation of variability in data. Therefore, quality control (QC) is of high importance, especially in the normative modeling setting. While manual QC on massive datasets is costly and not practical, there is still no bullet-proof automatic QC method available in the field. For this study, based on recent studies in this area, we used FreeSurfer’s Euler number (that summarizes the topological complexity of the reconstructed cortical surface) as a criterion for the quality of scans, which shows a good correspondence with manual ratings of scan quality (Rosen et al., 2018; Sánchez et al., 2021). Even though our manual inspection shows its reasonable performance, but we see addressing the open problem of developing automated QC as a decisive step toward reliable normative modeling, and we recommend this be given careful attention in future applications.

## 5 Conclusions

In this study, we delineated the components involved in the life-cycle of normative modeling. We further elucidated the essential requirements of the normative modeling life-cycle to overcome the challenges imposed in the model development and deployment in real-world clinical applications. Then, we introduced a simple yet effective probabilistic federated learning approach to satisfy those requirements. The proposed hierarchical Bayesian regression method is quite flexible and accommodates a full range of parametric/non-parametric and linear/non-linear functions for modeling the signal mean and homoscedastic/heteroscedastic variance. On massive experimental data and in realistic scenarios, the HBR showed superior performance in deriving normative ranges of cortical thicknesses compared to its alternatives. In the longer run, we believe our methodological contributions provide a significant step toward bringing precision medicine to the diagnosis and treatments of complex brain disorders.

## Acknowledgements

This work was supported by the Dutch Organisation for Scientific Research via Vernieuwingsimpuls VIDI fellowships to AM (016.156.415) and CB (864.12.003). The authors also gratefully acknowledge support from the Welcome Trust via digital Innovator (215698/Z/19/Z) and strategic awards (098369/Z/12/Z) and the Marie Sklodowska-Curie grant(895011).

## Supplementary Material

### Data after ComBat Harmonization

In pooling-after-harmonization models, we used ComBat for harmonizing data. The Python implementation at https://github.com/Warvito/neurocombat_sklearn is employed. Age and gender are used in the design matrix to ensure that their variability is preserved in data. Fig. S1 shows the distribution of the median of cortical thicknesses across 148 brain regions. Compared to Fig. 6 in the main text, it seems the scanner effect is successfully eliminated.

**Fig. S 1:**
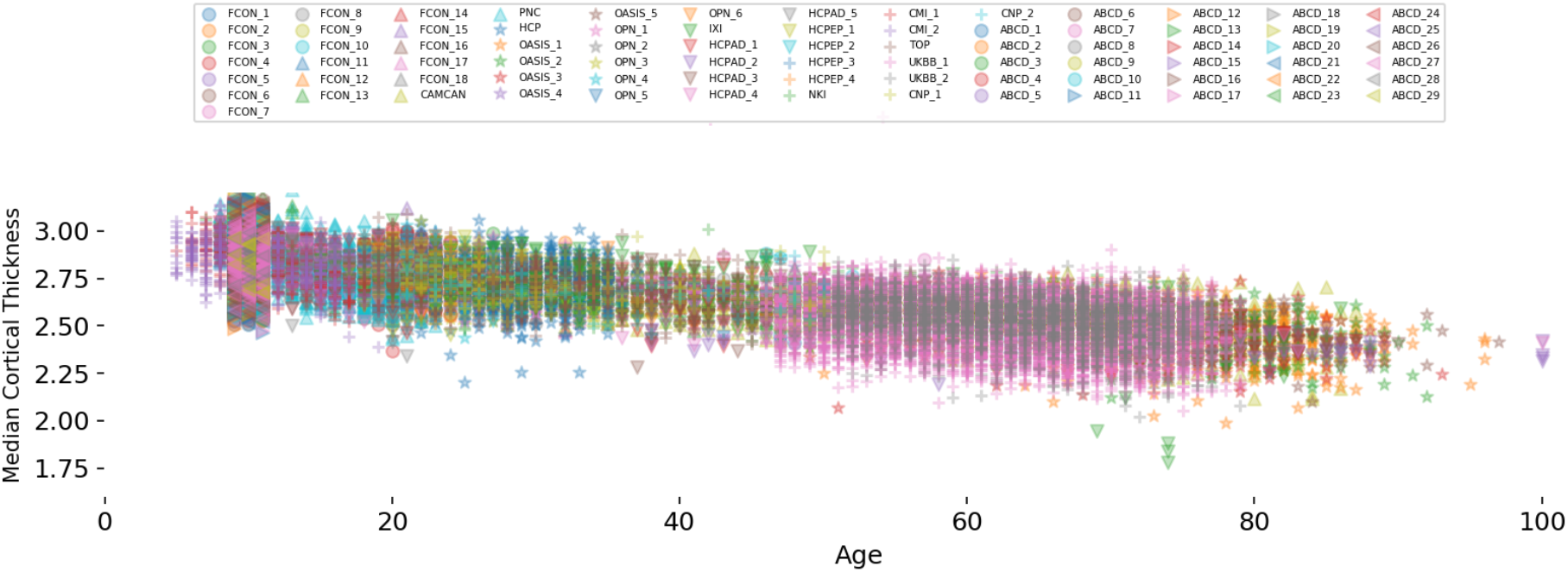
Median cortical thicknesses across 79 scanners after ComBat harmonization. The additive and multiplicative site-effects are successfully eliminated after harmonization.

### Simulation Study

In this simulation study, we aim to experimentally illustrate the theoretical limitations of data harmonization in a normative modeling setting. Following the reasoning in Sec. 2.3.2 and Fig. 2, we assume the signal **y** ∈ ℝ^*n*×1^ contains three components 1) the known source of variance that is measured via a covariate **x** ∈ ℝ ^*n*×1^; 2) the unknown variance **z** ∈ ℝ ^*n*×1^; and 3) the site-effect. In two exaggerated simulation scenarios, we assume that a hidden site-specific factor is present in the data. We use the following function to simulate samples from two sites:

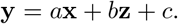

In the first scenario, we assume the hidden factor affects the variability of signal. Thus we set *a* = −0.4 and **x** ∈ [0, 10] as fixed for both sites (the effect of the measured covariate is fixed). We use *c* to simulated the additive site-effect and set it to 10 and 20 for *Site 1* and *Site 2*, respectively. On the other hand, we respectively set **z** to (**x** + 2)/3 and (12 − **x**)/3 for *Site 1* and *Site 2*, and assume 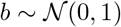. This results in different heteroscedastic variations across two sites. The resulting simulated data are depicted in Fig. S2(a). We used naive pooling, pooling after data harmonization using ComBat, and HBR methods to model the signal with respect to the covariate. In all cases, a linear model with heteroscedastic noise is employed. The results are shown in Fig. S2(b-d). As expected, the quantiles estimated using the naive pooling approach are highly contaminated with the site-effect and are completely misrepresented. Therefore, the resulting z-scores are highly confounded with the site-effect. We further used a linear SVM classifier in 5-fold cross-validation to classify the z-scores into sites that results in 0.98 ± 0.02 balanced accuracy in the naive pooling scenario. This is while the site-effect is not present in the resulting z-scores after harmonization and using the HBR method (0.42 ± 0.07 and 0.48 ± 0.07 respectively). However, the estimated quantiles in the case of harmonized data are misleading. For example, an outlier sample (marked by a red cross) lies in the 25% quantile. This is while in the HBR case, the quantiles are estimated on original data and separately for two sites. Thus, the outlier sample remains outside the 1% centile.

In the second scenario, we assume the hidden factor affects the slope of the effect. In this case, we set **z** to 1.5 for both sites. Instead we use different slopes for two sites setting *a* to 1 and −1 for *Site 1* and *Site 2*, respectively. The rest of the parameters are set as they were in the first simulation. Here, the hidden factor correlates with both covariate and site-effect. The resulting simulated data are shown in Fig. S3(a). The results of normative modeling using three different methods are shown in Fig. S3(b-d). In this case, since the hidden factor correlates with the known covariate, the ComBat is completely disarmed, thus, the harmonization has almost no effect on the data. Thus, the quantiles estimated using both pooling approaches (with and without harmonization) are completely misrepresented. This is while the HBR provides a reasonably good estimate of quantiles for two sites.

**Fig. S 2:**
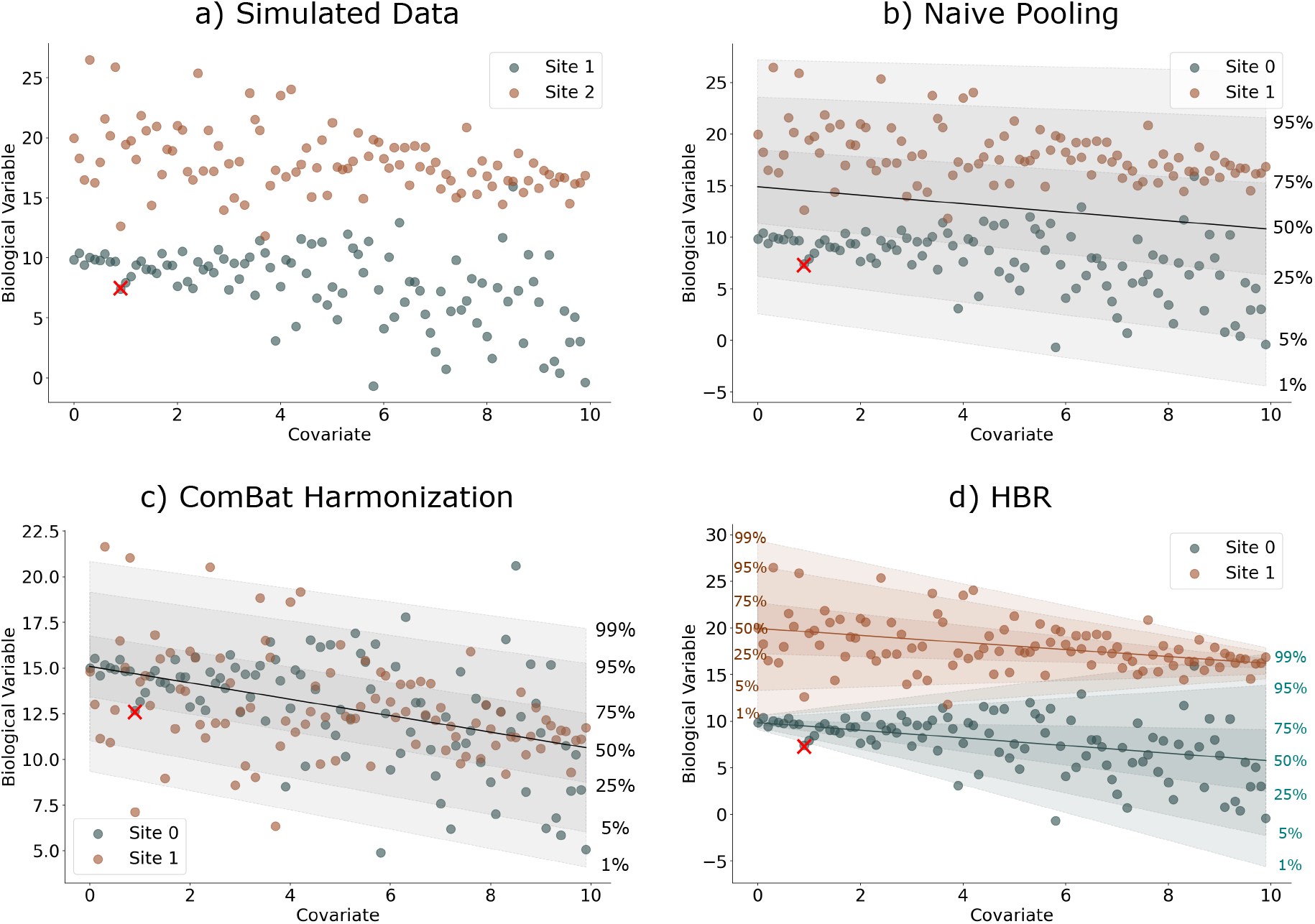
a) The simulated data in the first simulation scenario in which a site-specific hidden factor affects the variability of the signal. The normative ranges are estimated using b) naive pooling, c) pooling after ComBat harmonization, and d) HBR. The estimated quantiles are misrepresented in the naive pooling case due to the site-effect. Even though the site-effect is removed after harmonization, the estimated quantiles are still misrepresented, and an outlier sample (marked by a red cross) remains within the normal range. This problem is solved when using HBR for normative modeling.

Our simulations demonstrate how using data harmonization before normative modeling results in misleading quantile estimation, especially when a hidden variable correlates with site-effect and the covariate of the interest. Our analysis also highlights the differences between HBR and harmonization with ComBat. Specifically, ComBat removes the estimated location and scale parameters for each site from the data. In contrast, HBR estimates a *separate* location, slope, and scale for each site but does not remove them from data. This allows the model to capture different effects across sites and can accommodate unmeasured sources of variance. This feature could be useful, for example, in a lifespan study where site variation is highly correlated with age (as it is usually the case) and where there is heteroscedastic variance across the lifespan (*e.g.*, higher variance in older age ranges).

### HBR with Heteroscedastic Noise Model

The proposed HBR framework is capable of modeling heteroscedasticity. To evaluate the effect of accounting for heteroscedastic noise in modeling our experimental data, we conducted an experiment to compare the homoscedastic and heteroscedastic models. We have used different parametrization for parameters including linear, polynomial, and B-spline. Fig. S4 summarizes the results. Given the characteristics of experimental data, using heteroscedastic noise models for the variance provide a little (in the linear case) or no improvement over the homoscedastic models while having higher model complexity.

**Fig. S 3:**
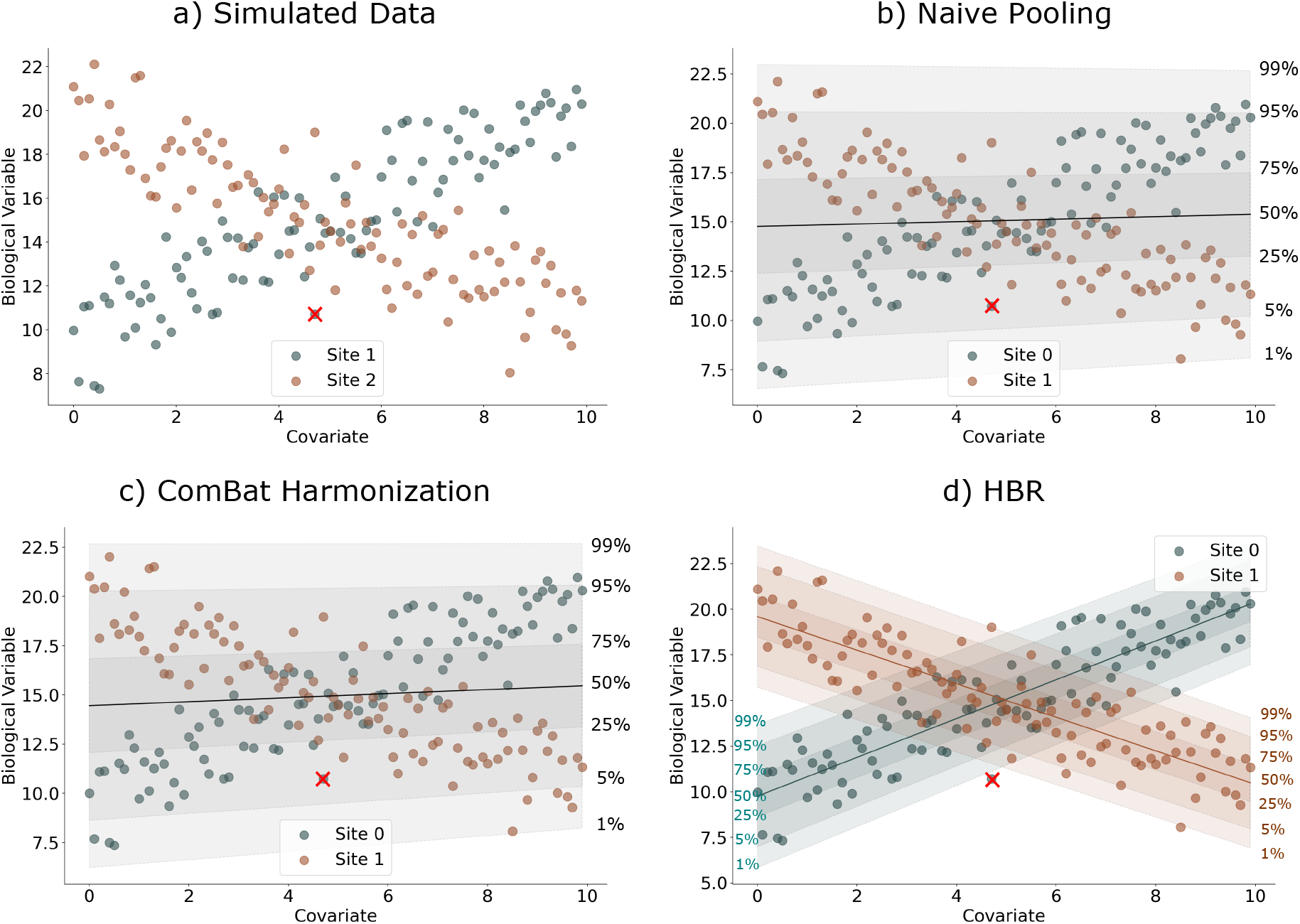
a) The simulated data in the second simulation scenario in which a site-specific hidden factor affects the slope of the signal. The normative ranges are estimated using b) naive pooling, c) pooling after ComBat harmonization, and d) HBR. The estimated quantiles are misrepresented in both pooling scenarios because the ComBat fails in removing the site-effect. Thus, an outlier sample (marked by a red cross) is read within the normal range. This problem is solved when using the HBR for normative modeling.

**Fig. S 4:**
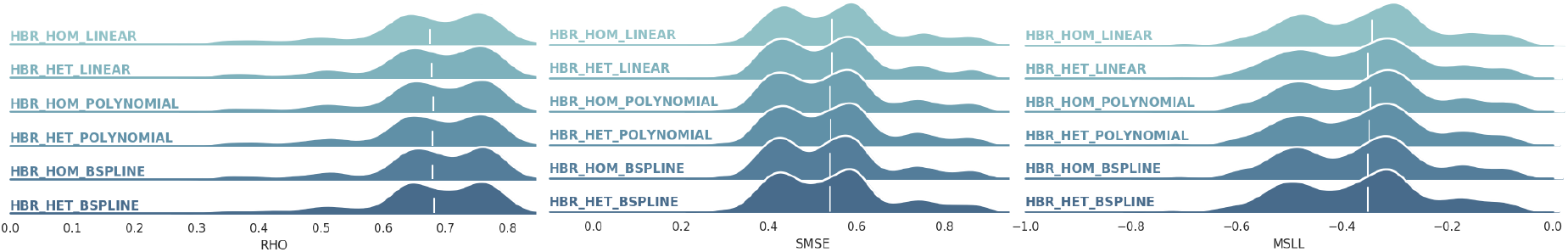
Comparison between regression performance of HBRs with homoscedastic and heteroscedastic variance models. On these experimental data, using more complex heteroscedastic models provides little or no improvement over simpler homoscedastic models.

